# Decision-making processes in perceptual learning depend on effectors

**DOI:** 10.1101/2022.06.29.498152

**Authors:** Vladyslav Ivanov, Giorgio L. Manenti, Sandrin S. Plewe, Igor Kagan, Caspar M. Schwiedrzik

## Abstract

Visual perceptual learning is traditionally thought to arise in visual cortex. However, typical perceptual learning tasks also involve systematic mapping of visual information onto motor actions. Because the motor system contains both effector-specific and effector-unspecific representations, the question arises whether visual perceptual learning is effector-specific itself, or not. Here, we study this question in an orientation discrimination task. Subjects learn to indicate their choices either with joystick movements or with manual reaches. After training, we challenge them to perform the same task with eye movements. We dissect the decision-making process using the drift diffusion model. We find that learning effects on the rate of evidence accumulation depend on effectors, albeit not fully. This suggests that during perceptual learning, visual information is mapped onto effector-specific integrators. Overlap of the populations of neurons encoding motor plans for these effectors may explain partial generalization. Taken together, visual perceptual learning is not limited to visual cortex, but also affects sensorimotor mapping at the interface of visual processing and decision making.

## Introduction

A core characteristic of the brain is its ability to learn. Learning is not limited to acquiring new motor skills or a foreign language – visual perception is also highly plastic. Training enables us to improve our visual abilities well into the hyperacuity range [1] and to learn to see what is initially invisible [2]. Such training effects are known as visual perceptual learning (VPL). VPL occurs not only during critical periods of development, but also in adult brains. This has profound implications for how we think about brain organization and function, yet little is known about the principles and neural mechanisms underlying adult VPL.

Classical theories suggest that VPL occurs in visual cortex [3-5]. This idea rests upon the striking correspondence between the known selectivities of neurons in early visual areas and the selectivity of behavioral learning effects that are often found in VPL studies. For example, VPL has been found to be specific to the orientation, location [6], polarity [7], and even eye of origin [8] of the training material, resembling the tuning properties of V1 neurons. Accordingly, many classical theories argue that VPL increases the selectivity of neurons in early sensory areas.

While VPL-induced increased selectivity of neurons in early visual cortex has indeed been observed through single cell recordings in monkeys [reviewed in 9], results are somewhat inconsistent across studies [10], the observed changes in selectivity are too small to explain the behavioral benefits [11], and modelling points to limitations on how much information can be gained by sharpening tuning curves [12]. These results suggest that sharpening of neural tuning in visual cortex is only one of the possible mechanisms of VPL.

More recent studies have started to investigate a previously neglected aspect of VPL, sensorimotor mapping. In a typical VPL task, two stimulus alternatives, e.g., clockwise or counterclockwise orientation of lines, have to be mapped onto two specific motor actions, e.g., a leftward or rightward saccade. This implies that visual information needs to be brought together with motor plans that correspond to the trainee’s choice. This suggests selective strengthening of connections between visual and motor neurons as another candidate mechanism of VPL. Indeed, a groundbreaking study found that VPL changes the activity of neurons in the lateral intraparietal area (LIP), an area in posterior parietal cortex that is involved in oculomotor decision-making and where sensorimotor mappings can take place [13]. This resonates with theories that frame VPL as a change in the readout of visual information by decision-making areas [14]. A role of motor areas in VPL is further suggested by several behavioral findings, e.g., that motor tasks improve VPL consolidation [15], and, reversely, that VPL can improve oculomotor performance [16].

If VPL affects sensory-motor mapping between visual cortex and the motor system, this would suggest another form of specificity in VPL, namely effector specificity. This is because the motor system contains neurons that code motor plans for specific effectors. These are highly prevalent in the primary motor cortex, but also in parietal cortex: many neurons in the so-called parietal reach region (PRR) code for arm movements, while many neurons in LIP are saccade specific [17-20]. During VPL, connections between stimulus-specific visual neurons and effector-specific motor neurons could thus be strengthened, resulting in stimulus- and effector-specific behavioral learning effects. However, areas like LIP also encode components that are effector-independent [18,21-24]. This raises the possibility that VPL is effector-independent, too, even if sensorimotor mapping is involved.

**Figure 1:**
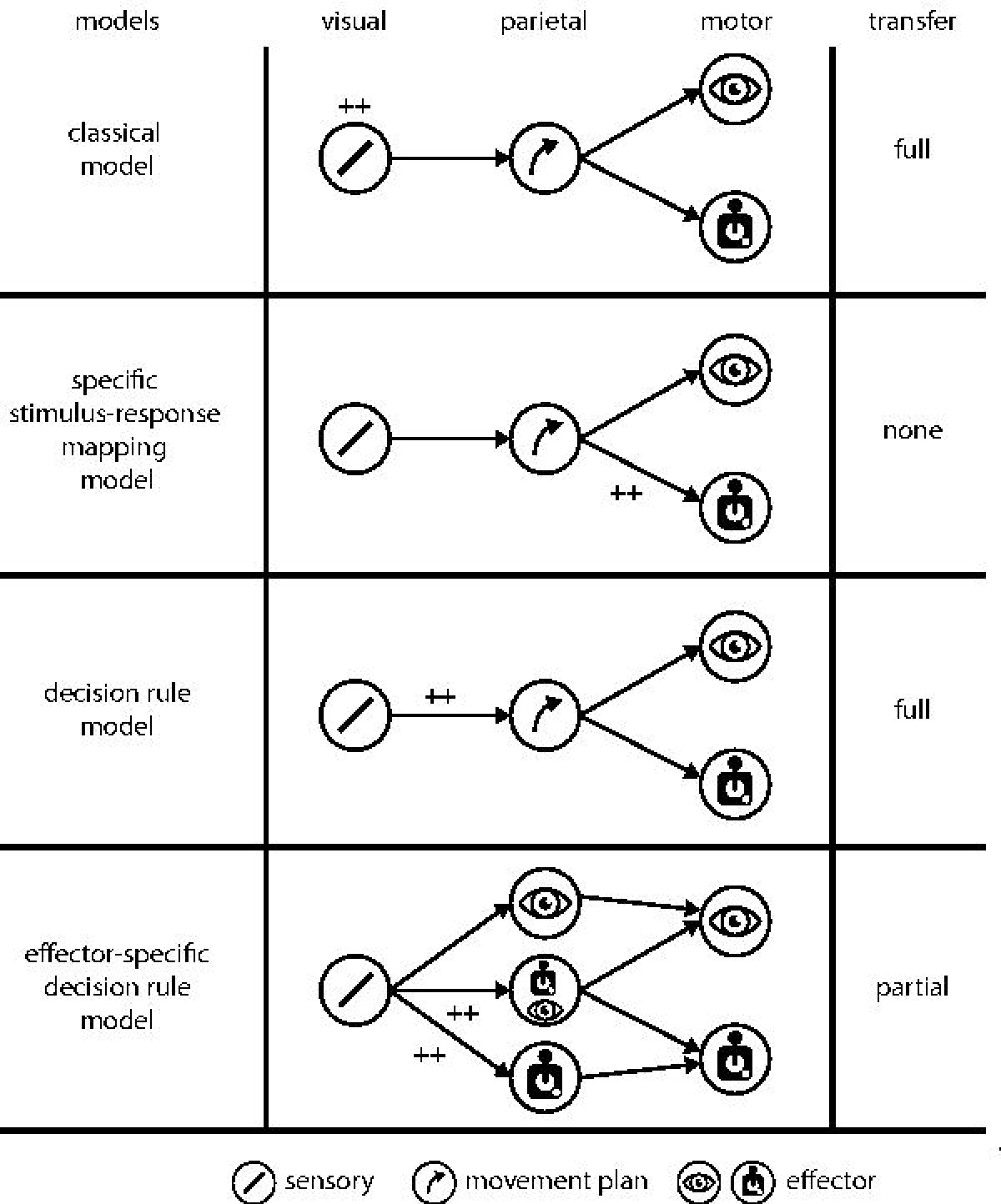
Possible models for effector specificity in visual perceptual learning. Classical model: learning occurs in visual cortex and results in stimulus-but not effector-specific learning effects. Changing the effector after training results in full transfer of learning effects to the new effector. Specific stimulus-response mapping model: visual information is mapped onto effector-specific motor neurons. This would result in stimulus- and effector-specific learning effects. Decision rule model: stimulus-response mapping takes place between visual neurons and effector-independent neurons involved in decision making and movement planning. This results in stimulus-specific learning effects that generalize across effectors. Effector-specific decision rule model: Sensorimotor mapping occurs between visual neurons and effector-specific integrators. Overlap between effectors at the integration stage results in partial transfer between the trained and untrained effectors.

A number of studies have started to investigate effector-specificity of VPL, but results are mixed, leaving open several possible models on how sensory and/or motor circuits are affected by training (Fig. 1): in the above-mentioned *classical model* of VPL, learning occurs in visual cortex and results in stimulus-but not effector-specific learning effects. Hence, changing the effector after training would result in full transfer of learning effects to the new effector. Studies that report effector-independent VPL effects can be interpreted this way [25]. In the *specific stimulus-response mapping model*, visual information is mapped onto effector-specific motor neurons in motor cortex [26], parietal cortex [18,19], or subcortical structures [27]. This would result in stimulus- and effector-specific learning effects. Evidence for high effector-specificity VPL has been reported in some studies [28,29]. However, these experiments often also involved a change in the task (e.g., binary categorization versus continuous adjustment), which possibly requires different readout weights. Hence, these results could also be interpreted to reflect the well-known task-specificity of VPL [30] instead of effector-specificity. Most recently, a *decision rule model* has been put forward [25]. Here, stimulus-response mapping takes place between visual neurons and effector-independent neurons involved in abstract movement planning and decision making in parietal cortex or subcortical structures like the pulvinar [31]. This would result in stimulus-specific learning effects that generalize well across effectors. This has indeed been reported [25].

The studies investigating effector-specificity in VPL have so far exclusively focused on accuracy as a measure of learning and transfer. However, this only considers the end result of a decision-making process and disregards subcomponents involved in the full sensorimotor processing arc. Here, we use a model-based approach, the well-established drift diffusion model (DDM) of decision making [32], to dissociate different aspects of this arc and learning effects upon them. The DDM relates accuracy to reaction times and allows investigation of the rate of evidence accumulation (the drift rate parameter), the decision bound (i.e., when a decision is reached), non-decision time (related to encoding of sensory information and execution of motor actions), and decision bias for one or the other stimulus alternative. This approach has previously been shown to reveal subcomponents of VPL [33-36], but has so far not been applied to the question of effector-specificity in VPL.

We train subjects in an orientation discrimination task. We focus on orientation VPL since stimulus-specific VPL effects are well documented for this feature [6]. Subjects practice for several days to discriminate orientations and to indicate their choice with a joystick movement (Experiment 1). After the last training session, we require them to carry out the same task but to indicate their choice with an eye movement. Our transfer task thus differs from training only in the effectors, allowing us to dissociate the above-mentioned models. In Experiment 2, we train subjects to indicate their choice with a pointing movement instead that closely matches the spatial transformations required for directing saccades during the transfer task. This allows us to further differentiate whether any possible specificities are due to differences in spatial transformations or the effectors themselves.

To preview, across the two experiments, we find that VPL depends on the effector. The DDM taking reaction times into account shows that the specificity arises in the rate of sensory evidence accumulation (i.e., the drift rate), an integral component of decision-making. This is most compatible with a new model, where visual information is mapped onto effector-specific integrators.

## Methods

### Participants

A total of 44 healthy human volunteers (28 female, 1 left-handed, mean age 25.61 yrs, *SD* 6.94 yrs) participated in this study. All subjects had normal or corrected-to-normal vision, reported no history of neurological or psychiatric disease, and gave written informed consent before participation. No sample size estimate was performed, but sample size was selected based on previous studies. Because most comparisons were within subjects, we used convenience sampling. 22 subjects (12 female, 1 left-handed, mean age 23.5 yrs, *SD* 2.32 yrs) participated in Experiment 1 involving joystick and eye movements. 3 subjects had to be excluded from data analysis because lack of significant learning (final *n*=19, 12 female, 1 left-handed, mean age 23.68 yrs, *SD* 2.4 yrs). 12 subjects participated in Experiment 2 involving reaches on a touchscreen and eye movements (*n*=12, 10 female, mean age 26.8 yrs, *SD* 11.78 yrs). 2 subjects were excluded from data analysis because lack of significant learning (final *n*=10, 9 female, mean age 25.8 yrs, *SD* 1.86 yrs). 10 subjects participated in a control experiment for baseline differences between joystick and eye movements (*n*=10, 6 female, mean age 27.3 yrs, *SD* 6.21 yrs). All procedures were in accordance with the Declaration of Helsinki and approved by the Ethics Committee of the University Medical Center Göttingen (protocol number 29/8/17).

### Stimuli and procedure

In all experiments, subjects were trained on an orientation discrimination task. Training took place over four to six consecutive days with one training session per day. On the day after the last training session, subjects were instructed to perform the same task, but with a different effector (see below). Stimuli were presented on a gamma-corrected LCD monitor (ViewPixx EEG, refresh rate 120 Hz, resolution 1920 × 1080 pixel, viewing distance 65 cm) in a darkened, sound-attenuating booth (Desone Modular Acoustics). Stimulus delivery and response collection were controlled using Psychtoolbox [37] running in Matlab (The Mathworks, Inc.) on Windows 10. During all experiments, we continuously acquired pupil and gaze measurements using a high-speed, video-based eye tracker (SR Research Eyelink 1000+). Data were sampled at 1000 Hz from both eyes. Subjects in Experiments 1 and 2 were paid €8 per hour. To assure constant motivation over the training sessions, subjects received a bonus of €2 if they improved by 10% from the previous training session. Subjects in the control experiment were paid 12€ per hour to assure a similarly high level of motivation as in Experiments 1 and 2.

**Figure 2:**
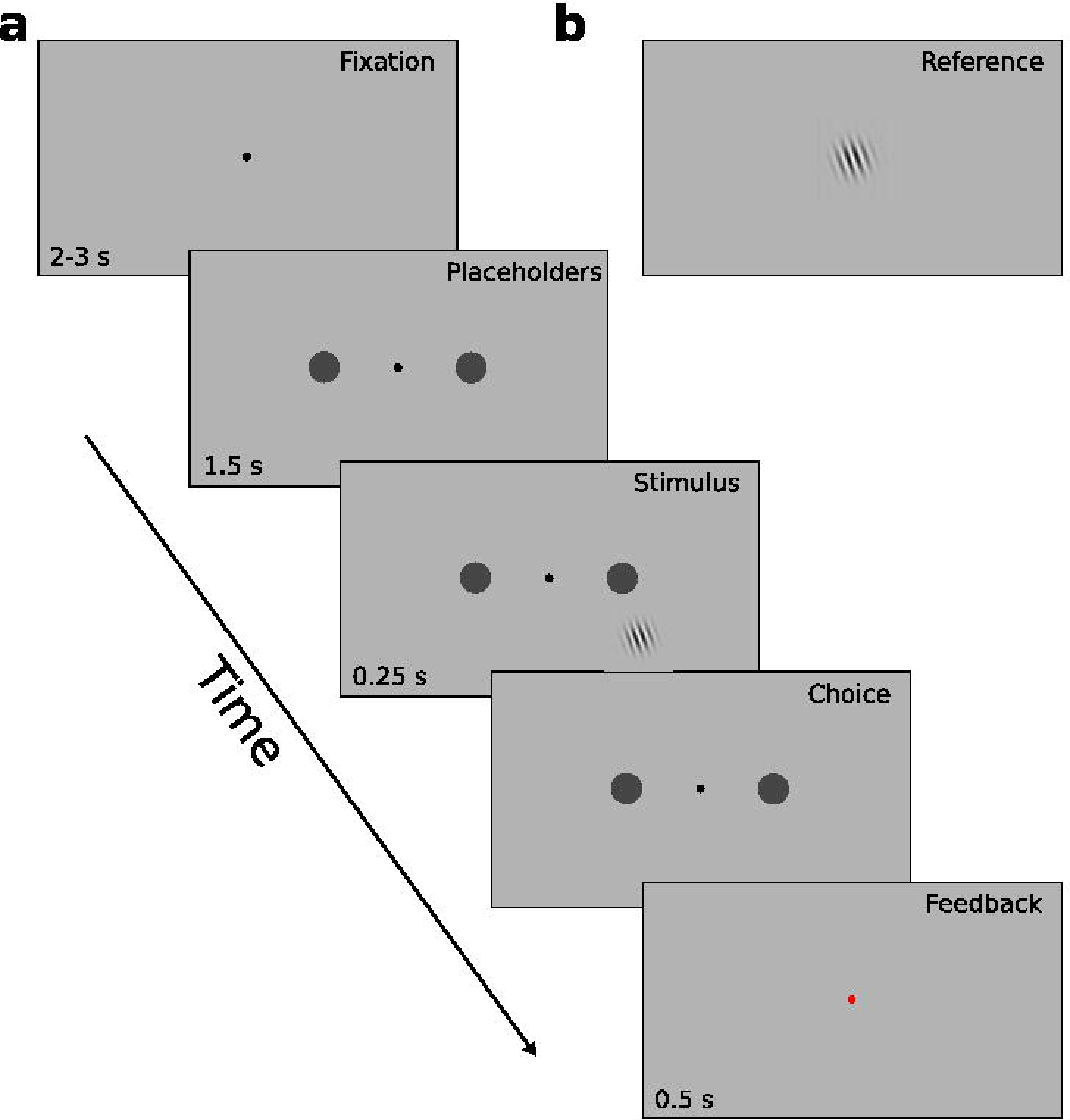
Design of Experiment 1. **A.** Each trial started with a fixation phase (2-3 s). Then, placeholders for the saccade version of the task appeared (1.5 s), followed by the presentation of the Gabor grating (0.25 s). Once the Gabor grating disappeared, subjects had 1 s to indicate their choice (with a joystick movement during the training phase or with a saccade during the transfer session). Feedback on accuracy was presented by changing the color of the fixation dot to red or green (0.5 s). **B.** The reference grating was shown for 2 s in the center of the screen before and in the middle of each block (in total 8 times per session).

#### Experiment 1

In Experiment 1, subjects were trained in orientation discrimination and responded with a joystick movement during training, while they had to respond with a saccade in the transfer session. Subjects had to decide whether a Gabor grating (size 2.4 dva, spatial frequency 3.3 cpd, luminance 43.4 cd/m^2^) was tilted clockwise or counterclockwise with respect to the reference stimulus. The reference stimulus was a Gabor grating with identical parameters tilted 20° counterclockwise from the vertical meridian presented at the center of gaze. The task stimuli were presented in six logarithmically spaced difficulty levels (±Δ = [0°, 0.31°, 0.56°, 1°, 1.77°, 3.16°]) with 20 trials for each of the orientations, respectively. The stimuli were presented in pseudo-random order at 8.65° eccentricity in the lower right quadrant against a grey background (39.5 cd/m^2^).

Each trial started with a fixation phase of 2-3 s duration (Fig. 2). Then, placeholders for the saccade version of the task appeared for 1.5 s at 8° eccentricity (distance from task stimulus 6.7 and 14.5 dva, respectively), followed by the presentation of the Gabor grating for 250 ms. Subjects then had 1 s to respond. The start of the response period was indicated by the disappearance of the Gabor grating. Feedback on accuracy was presented by changing the color of the fixation dot to red (incorrect) or green (correct) for 0.5 s. The next trial started 1.5 s later. If subjects did not respond within 1 s, a message “too late” appeared on the screen and the trial was repeated later during the block. Similarly, if subjects broke fixation throughout the trial, the trial was aborted and repeated at the end of the block.

Each training session included 220 trials evenly distributed among four blocks. Subjects were free to take short breaks between blocks. There were additional breaks in the middle of each block to display feedback about performance and reaction times. The reference was shown for 2 s in the center of the screen before and in the middle of each block (in total 8 times per session).

Subjects were instructed to respond as accurately as possible, however, they also had to consider the time limitation. To respond, subjects had to tilt a joystick (Logitech Extreme 3D Pro, either in standard configuration or in a customized configuration for left-handed subjects) with their dominant hand to the left if a stimulus was counterclockwise rotated and to the right if clockwise (while maintaining fixation). Hand dominance was assessed at the beginning of the experiment with the Edinburgh Handedness Inventory [38]. In the transfer session, subjects were instructed to instead direct their gaze to the left or right placeholder, respectively. Subjects thus had to learn an arbitrary mapping of a visual stimulus to a response alternative, and potentially remap this association to a new effector in the transfer session. Previous research has shown that effectors do not affect accuracy pre training [39-41]. We focus on effector-specificity since stimulus-specificity does not differentiate the different models for sensorimotor mapping in VPL.

Participant had to reach at least 75% correct responses before we transitioned to the transfer session. All but two subjects reached this performance level within four days; two subjects required five and six days of training, respectively.

**Figure 3:**
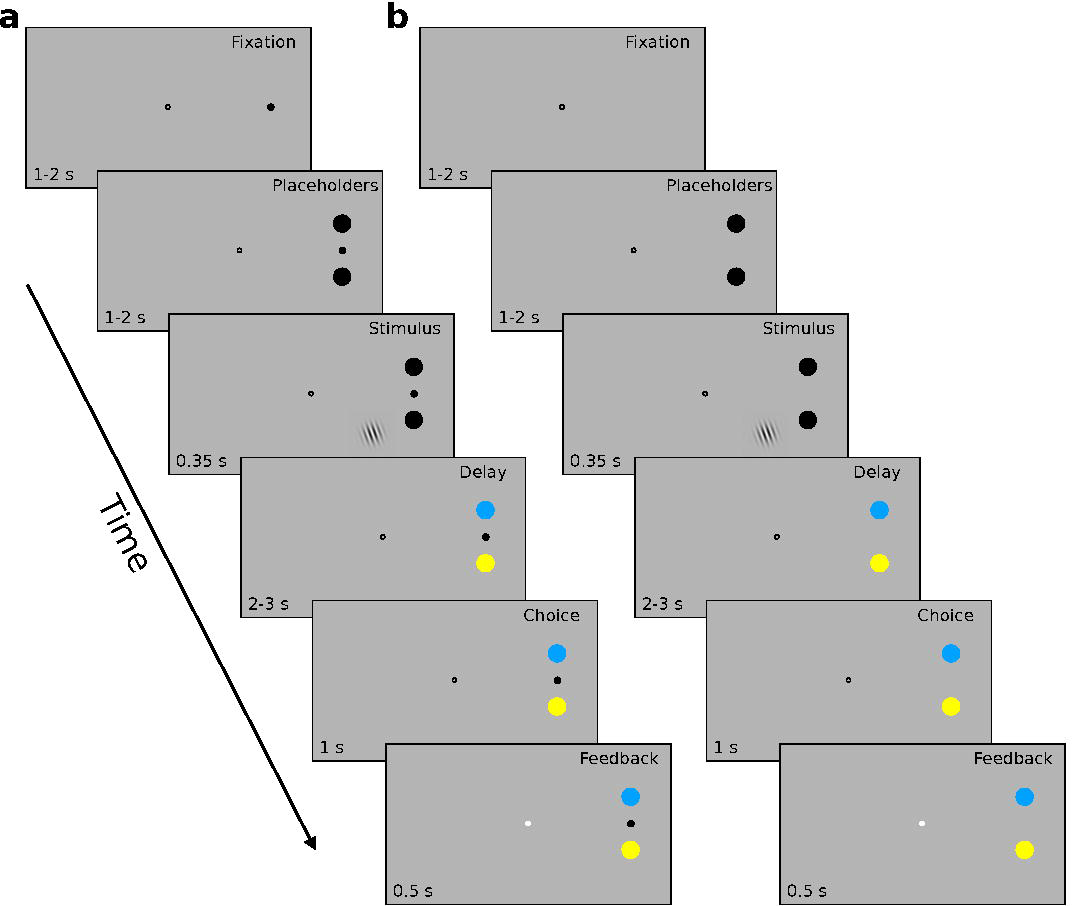
Design of Experiment 2. **A.** In the touchscreen version of the task, each trial started with a finger and eye fixation phase of (1-2 s). Subjects had to maintain central fixation with their gaze and simultaneously touched a finger fixation dot between the two target locations on the screen. Then, black placeholders appeared (1-2 s), followed by the presentation of the Gabor grating (0.35 s). A delay phase (2-3 s) started with a switch of the placeholders’ color to blue and yellow. Subjects had to respond within 1 s once the finger fixation dot disappeared. Feedback on accuracy was presented by changing the color of the fixation dot to blue or white (0.5 s). **B.** The trial structure of the saccade version of the task was identical to the touchscreen version, with the exception that the finger fixation dot was not shown. Instead, the disappearance of the central fixation dot indicated the start of the response period.

#### Experiment 2

In Experiment 2, subjects were trained on orientation discrimination and responded with a finger movement during training, while they had to respond with a saccade in the transfer session. We recorded finger movements by means of a transparent touch screen (Magic Touch, Tyco Touch Inc.), modelled after Pilacinsky and Lindner [42], which was mounted in front of the screen. Subjects were provided with an arm support (ErgoRest) and a finger stylus (TrueTip) to ensure ease of use.

As in Experiment 1, subjects had to decide whether a Gabor grating (size 2.4 dva, spatial frequency 3.3 cpd, luminance 10.9 cd/m^2^) was tilted clockwise or counterclockwise with respect to the reference stimulus. The reference stimulus was a Gabor grating with identical parameters tilted 20° counterclockwise from the vertical meridian. The task stimuli were presented in six logarithmically spaced difficulty levels (±Δ = [0°, 0.31°, 0.56°, 1°, 1.77°, 3.16°]) with 20 trials for each of the orientations, respectively. The stimuli were presented in a pseudo-random order at 8.65° eccentricity in the lower right quadrant against a dark grey background (6.4 cd/m^2^).

Each trial started with a finger and eye fixation phase of 1 to 2 s duration (Fig. 3). Subjects had to maintain central fixation with their gaze and simultaneously touched a finger fixation dot between the two target locations on the screen. Then, black placeholders appeared for 1 to 2 s at 11° eccentricity (distance from task stimulus 5 and 11.4 dva, respectively), followed by the presentation of the Gabor grating for 350 ms, a delay of 2 to 3 s during which the placeholders’ color switched from to blue and yellow, respectively. Subjects had 1 s to respond. The start of the response period was indicated by change of fixation dot. The next trial started 0.5 to 2 s later. If subjects did not respond within 1 s, the trial was repeated later during the block. Similarly, if subjects broke fixation throughout the trial, the trial was aborted and repeated at the end of the block. Feedback about correct and incorrect responses was provided with a blue or white dot, respectively, at fixation.

Each training session included 220 trials evenly distributed among four blocks. Subjects were free to take short breaks between blocks. The reference was shown for 2 s in the center of the screen before and in the middle of each block (in total 8 times per session).

Subjects were instructed to respond as accurately as possible. To respond, subjects had to move their finger from the finger fixation spot between the two targets to the blue target if the stimulus was tilted clockwise or to the yellow target if the stimulus was tilted counterclockwise (while maintaining central fixation). Hand dominance was assessed at the beginning of the experiment with the Edinburgh Handedness Inventory [38]. In the transfer session, subjects were instructed to instead direct their gaze to the blue or yellow placeholder, respectively. For all subjects, training lasted four days.

#### Control experiment for differences between effectors

To determine whether any differences between effectors were the result of training or already existed before training began, we conducted a control experiment comparing orientation discrimination performance between effectors. This was done in a separate group of subjects (*n*=10, 6 female, mean age 27.3 yrs, *SD* 6.21 yrs) to avoid possible effects of pretesting on generalization performance. The task was identical to Experiment 1, and subjects were asked to complete 220 trials with one effector and, after a break, 220 trials with the other effector. The order of effectors was counterbalanced across subjects.

### Analyses

The Signal Detection Theory parameter dill [43] was computed per subject and session using the loglinear correction [44]. *d’* were entered into repeated measures analyses of variance (rmANOVA) with factors session (4 levels for training session effects, or 2 levels for last training session to transfer effects), difficulty (5 levels), and their interaction. Reaction times were entered into a rmANOVA with factors session and correctness (levels correct, incorrect). Two subjects required more than four days of training (5 and 6) in Experiment 1; therefore, we restrict our analyses to the last four training sessions. Due to technical issues, we lost the data from the third training session for one subject in Experiment 1. Thus, to be able to compute rmANOVA across training sessions (1-4), we excluded the subject (total *n*=18) from these analyses, but we included the subject in the transfer test (last training versus transfer, total *n*=19). For *t*-tests and rmANOVAs, we compute Hedges’ *g* and partial *η^2^*, respectively, as effect sizes, using the Measures of Effect Size Toolbox [45]. In addition, to quantify transfer performance, a Specificity Index [33] was calculated, as follows:

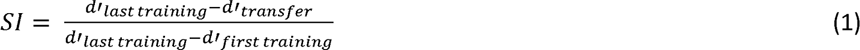

The index reflects the training session used to compute the median *d’*.

### Drift Diffusion Model

Sequential sampling models like the DDM assume that decision making is a stochastic process of evidence accumulation about a stimulus in time [32]. Decisions are made when the accumulation process reaches an evidence threshold (decision criterion) for one of the alternatives. The rate of accumulation and the decision criteria determine the reaction time (RT), and fluctuations in the process are the reason for errors and the distribution of RTs. The DDM can account for effects of different experimental conditions from accuracy, mean RT for correct and error responses, and the shape of RT distributions for correct and error responses. The full DDM includes seven parameters: drift rate (*v*) and its trial-by-trial variability (*η*), non-decision time (*T_er_*) and its trial by-trial variability (*s_t_*), boundary separation (*a*), the starting point of the decision process (*z*) and its trial-by-trial variability (*s_z_*). *v* is the key component of the model, as it describes the rate of evidence accumulation and positively correlates with the accuracy level of subjects. *a* is the distance between decision criteria for both alternatives, and the bigger the distance is, the longer it takes to make a decision. The evidence accumulation starts from *z*, which lies between 0 and *a*. *z* can be positioned closer to one of the alternatives, which biases the decision process toward it. *T_er_* accounts for the non-decision phases: stimulus encoding and response output. The DDM states that *v* and *T_er_* are independent, and hence, that manipulations of the effector only affect *T_er_* [46]. This assumption has been previously empirically validated [46], and it is thus common in the field to compare DDM parameters directly between effectors [e.g., 47]. Here, we follow the same logic.

We use the Hierarchical Drift Diffusion Model (HDDM 0.8.0) [48], available as a toolbox in Python, to fit the model parameters to our data of Experiment 1. The HDDM employs Bayesian inference to recover the model parameters. First, the HDDM estimates the parameter values for a population (*p*_group_), and then it uses the population fit to constrain the parameter values for individual subjects (*p*_subj_). The hierarchical Bayesian approach performs significantly better than using *χ^2^* as the optimization criterion when data per subject is limited [49].

To identify the combination of model parameters which best explains our data, we fit several models built on theoretically meaningful parameter combinations (see Results) and compare them on the basis of the Deviance Information Criterion [DIC, 50].

In the best performing model, we estimate parameters for all sessions jointly while allowing *v* to vary across the sessions and the difficulty levels. The stimuli are encoded by their orientation (clockwise/counterclockwise), and *v i*s estimated for both orientations together. We allow *a*, *T_er_* and *z* to vary only between sessions because a priori, decision boundaries, encoding, muscle response time, and the orientation bias should be the same across difficulty levels when the latter are randomly intermixed. However, it has been argued that subjects can learn to adjust their decision bounds to difficulty levels [51,52]. We thus also explored a model in which decision bounds could vary across sessions and difficulty levels, which we report in the supplementary material. We did not include noise parameters capturing inter-trial variability in the drift-rate, the non-decision time and the starting point into any our models; for proper estimation of these parameters, the HDDM requires noticeably more samples per condition than we have available. We run the fitting algorithm four times with 30000 Markow-Chain-Monte-Carlo samples, 10000 samples burn-in, and discarding every 5^th^ sample.

We utilize two formal diagnostics to assess the convergence of the model: the Gelman-Rubin and Geweke method, respectively. The Gelman-Rubin method compares between-chain and within-chain variances for each model parameter [53]. Significant differences (diagnostics value >1.2 or 1.1 under strict conditions) between these variances indicate non-convergence [54]. We compute the diagnostic with the help of the kabuki (0.6.3) Python library. The Geweke method evaluates only within-chain fluctuation [55]. It calculates *z*-scores of the difference between the initial and subsequent segments of the chain. If the chain has converged, most points should be within two standard deviations of zero (*z*-score ∈ [-2, 2]). We chose the first 10% of the chain as the initial segment, and the last 50% divided into 20 segments. To compute the Geweke diagnostic, we use the PyMC (2.3.8) Python library.

To compare model parameters between conditions within subjects, we assess the overlap of the respective posterior distribution. To assess differences between groups, we compare the distribution of differences between conditions between groups and report the probability of one group having a higher estimated difference in parameter value than the other group [56].

## Results

### Learning and transfer effects on accuracy and reaction time

We find that across all five difficulty levels, subjects in Experiment 1 significantly improve their performance (*d’*) as function of training, with increases ranging between 0.60 and 1.93 (Fig. 4A, mean increase between first and last training session 1.25, *t*(18)=-11.78, *p*<0.001, *g*=3.04; rmANOVA main effect of session, *F*(3, 51)=138.94, *p*<0.001, *η*^2^_*p*_ =0.519). Subjects learn more on easy than on difficult orientation differences (Fig. S1, main effect of difficulty, *F*(4, 68)=96.18, *p*<0.001, *η*^2^_*p*_ =0.647, interaction session × difficulty *F*(12, 204)=13.18, *p*<0.001, *η*^2^_*p*_ =0.145). When the effector is changed from a manual joystick response to a saccadic response in the transfer session, we find a significant decrease in performance relative to the last session across all difficulty levels (mean decrease 0.32, *t*(18)=6.41, *p*<0.001, *g*=0.98; rmANOVA main effect of session, *F*(1,18)= 21.48, *p*<0.001, *η*^2^_*p*_ =0.54 main effect of difficulty, *F*(4,72)= 290.68, *p*<0.001, *η*^2^_*p*_ =0.94, interaction session × difficulty *F*(4,72)=0.487, *p*=0.745, *η*^2^_*p*_ =0.02). This is also evident when we calculate the SI, which has a median of 0.30 (Fig. 4C, *p*<0.001, sign test, one-sided). In contrast, there are no differences in *d’* when the task is carried out with joystick or saccades in the control group in the absence of training (mean difference in *d’* 0.14, *t*(9)=1.10, *p*=0.653, *g*=0.21). Accordingly, the difference in *d’* between effectors in the trained group is significantly larger than the difference in the control group (mixed ANOVA, main effect of experimental group, *F*(1,27)=8.593, *p*=0.007, *η*^2^_*p*_ =0.241, no main effect of difficulty, *F*(4,108)=1.126, *p*=0.348, *η*^2^_*p*_ =0.04, no interaction experiment × difficulty *F*(4,108)=0.96, *p*=0.432, *η*^2^_*p*_ =0.03). This suggests that the drop in d’ we find in Experiment 1 when effectors change is learning-induced and cannot be explained by preexisting differences.

Mean reaction times for correct and incorrect responses do not differ from each other throughout the sessions and do not change significantly as a function of learning (all *p*>0.29, all *η*^2^_*p*_ < 0.11). When the effector is changed, reaction times become uniformly shorter (Fig. 4B and Fig. S1, mean difference 0.143 s, main effect of session, *F*(1,18)=91.56, *p*<0.001, *η*^2^_*p*_ = 0.83). As there was no significant main effect of correctness or any significant interaction (all *p*>0.09, all *η*^2^_*p*_ < 0.14) in reaction times, this is unlikely to constitute a learning effect and can be attributed to generally shorter latencies of saccades as compared to hand movements [40] that we also find in our control group (mean difference 0.12 s, rmANOVA main effect of effector, *F*(1,9)=6.731, *p*=0.029, *η*^2^_*p*_ = 0.428, all other effects *p*>0.84)).

**Figure 4:**
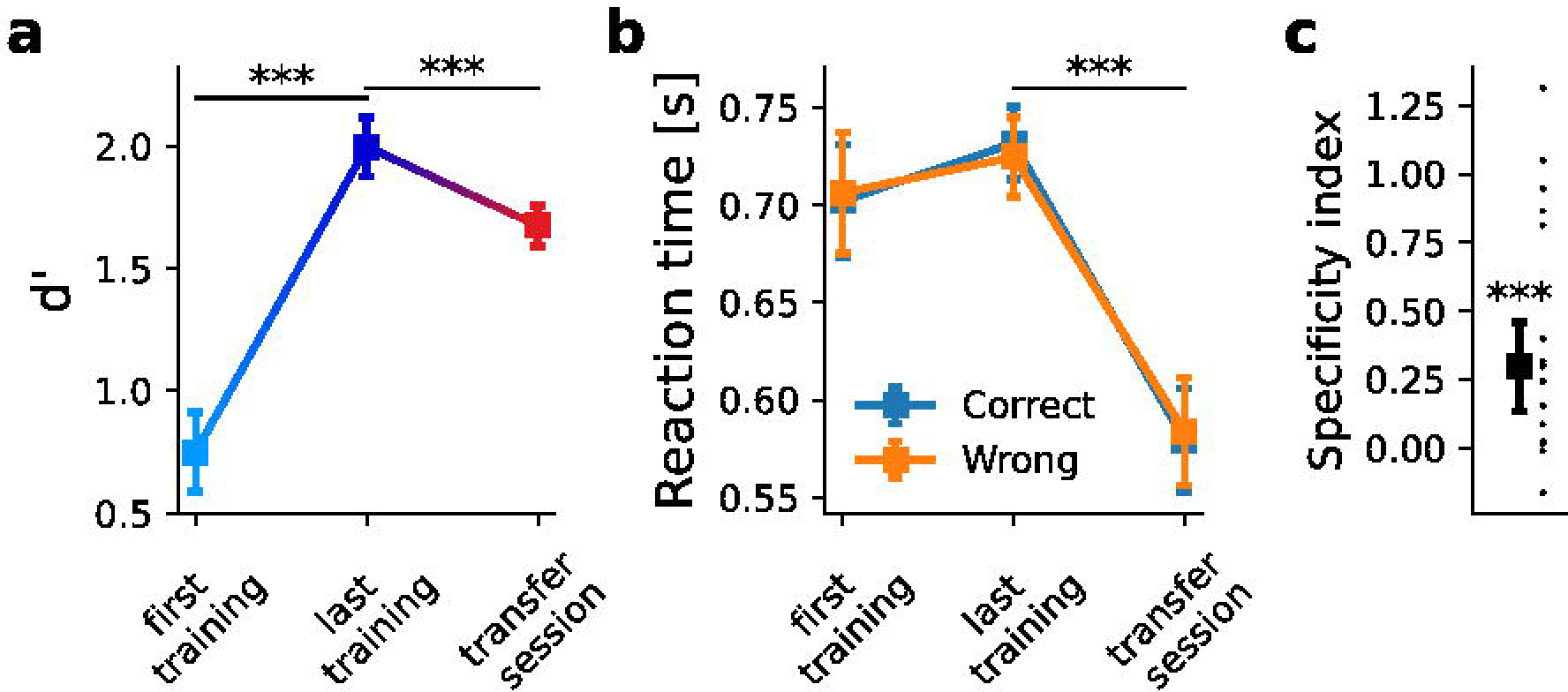
Accuracy and reaction times for Experiment 1. **A.** Mean accuracy (in units of d’) increases with learning from the first to the last training session (mean difference 1.25, t(18)=-11.78, p<0.001, g=3.04, two-sided) and drops from the level reached in the last training session when the effector is changed from a manual joystick response to a saccadic response (mean difference 0.32, t(18)=6.41, p<0.001, g=0.98, two-sided). **B.** Mean reaction times for correct and incorrect responses do not differ from each other throughout the sessions and do not change significantly as a function of learning (all p>0.29, all *η*^2^_*p*_ < 0.11). When the effector is changed, reaction times become uniformly shorter (main effect of session, F(1,18)=91.56, p<0.001, *η*^2^_*p*_ = 0.83, no main effect of correctness or interaction, all p>0.09, all *η*^2^_*p*_ < 0.14). **C.** Learning depends on the effector, as evidenced by a median Specificity Index of 0.30, which is significantly bigger than 0 (p<0.001, sign test, one-sided). Dots reflect individual subjects’ Specificity Indices. 17 out of 19 subjects have Specificity Indices >0. In panels A and B, error bars reflect the SEM, corrected for between-subject variability [80,81]. In panel C, error bars reflect the standard error of the median [82]. *** stand for p < 0.001, ** for p < 0.01, and * for p<0.05.

We find very similar results in Experiment 2, where subjects perform the same task with a precise reach on a touch screen. In this smaller sample (*n*=10), *d’* also improves with training over sessions (Fig. 5A, mean increase between first and last training session 0.82, *t*(9)=7.76, *p*<0.001, *g*=1.65; rmANOVA main effect of session, *F*(3,27)=24.46, *p<*0.001, *η*^2^_*p*_ = 0.20), with larger improvements for easier orientation differences than for difficult orientation differences (Fig. S1, main effect of difficulty, *F*(4,36)=106.65, *p*<0.001, *η*^2^_*p*_ =0.71, interaction session × difficulty *F*(12,108)=4.88, *p<*0.001, *η*^2^_*p*_ =0.12). When we change the motor component of the task from reaches to saccades in the transfer session, we again find a drop in discrimination across difficulty levels, albeit not statistically significant (mean decrease 0.12, *t*(9)=0.99 *p*=0.347, *g*=0.2; rmANOVA main effect of session, *F*(1,9)=0.98, *p*=0.347, *η*^2^_*p*_ =0.01, main effect of difficulty, *F*(4,36)=66.24, *p*<0.001, *η*^2^_*p*_ =0.69, interaction session × difficulty *F*(4,36)=1.97, *p*=0.119, *η*^2^_*p*_ =0.03). Specificity Indices nevertheless show that training is specific to the effector in 7 out of 10 subjects (Specificity Index >0); expressed in median SI across all subjects, this is 0.49 (Fig. 5C, *p*=0.035, sign test, one-sided).

**Figure 5:**
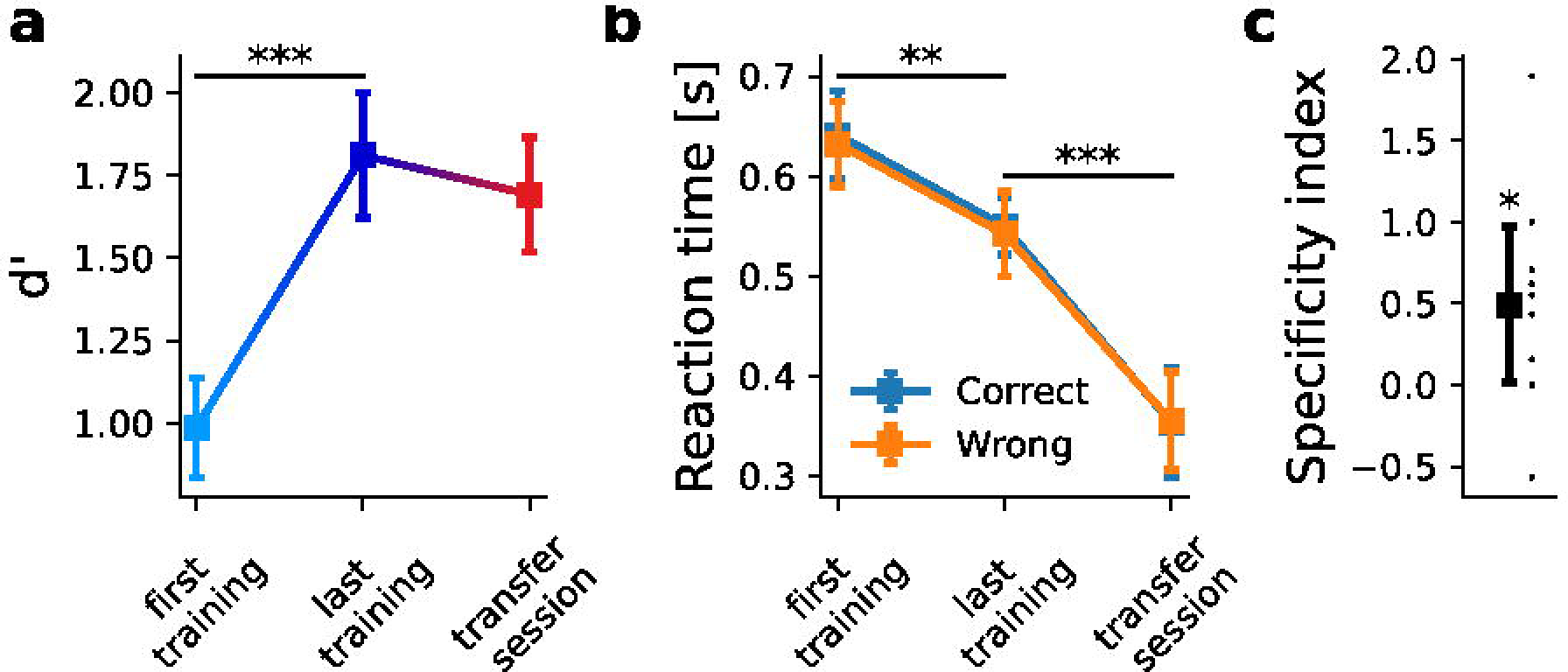
Accuracy and reaction times for Experiment 2. **A.** Mean accuracy (in units of d’) increases with learning from the first to the last training session (mean difference 0.82, t(9)=7.76, p<0.001, g=1.65, two-sided). There is a numerical, but no statistically significant drop in accuracy when the effector is changed from a manual joystick response to a saccadic response (mean difference 0.12, t(9)=0.99, p=0.345, g=0.2, two-sided). **B.** Mean reaction times decrease over sessions (mean decrease 0.09 s, t(9)=3.83, p=0.004, g=-0.97) and further drop significantly when the effector is changed (mean decrease 0.19 s, t(9)=-64.1497, p<0.001; g=-2.6325). **C.** Training is specific to the effector in 7 out of 10 subjects (Specificity Index >0), with a median Specificity Index of 0.49, which is statistically significantly bigger than 0 (p=0.035, sign test, one-sided). Dots reflect individual subjects’ Specificity Indices. In panels A and B, error bars reflect the SEM, corrected for between-subject variability [80,81]. In panel C, error bars reflect the standard error of the median [82]. *** stand for p < 0.001, ** for p < 0.01, and * for p<0.05.

Average reaction times decrease over sessions (Fig. 5B and Fig. S1, mean decrease 0.09 s, *t*(9)=3.83, *p*=0.004, *g*=-0.97; rmANOVA main effect of session, *F*(3,27)=10.94, *p*<0.001, *η*^2^_*p*_ =0.170, main effect of correctness, *F*(1,9)=1.20, *p*=0.301, *η*^2^_*p*_ =<0.001, interaction session × correctness *F*(3,27)=0.89, *p*=0.45, *η*^2^_*p*_<0.001) and further drop significantly when the effector is changed (mean decrease 0.19 s, *t*(9)=-6.14, *p<*0.001; *g*=-2.63; rmANOVA main effect of session, *F*(1,9)=37.74, *p<*0.001, *η*^2^_*p*_ = 0.673, main effect of correctness, *F*(1,9)=0.315, *p*=0.588, *η*^2^_*p*_<0.001, interaction session × correctness *F*(1,9)=1.80, *p*=0.213, *η*^2^_*p*_ =0.001). The latter can again be attributed to generally shorter reaction times for saccades than for reaches. Together, the SIs from Experiments 1 and 2 show that learning effects on accuracy in orientation discrimination depend on the effector without any change in the stimulus material or the task (median SI across both experiments=0.30, SE 0.13, *p*<0.001, sign test, one-sided).

**Figure 6:**
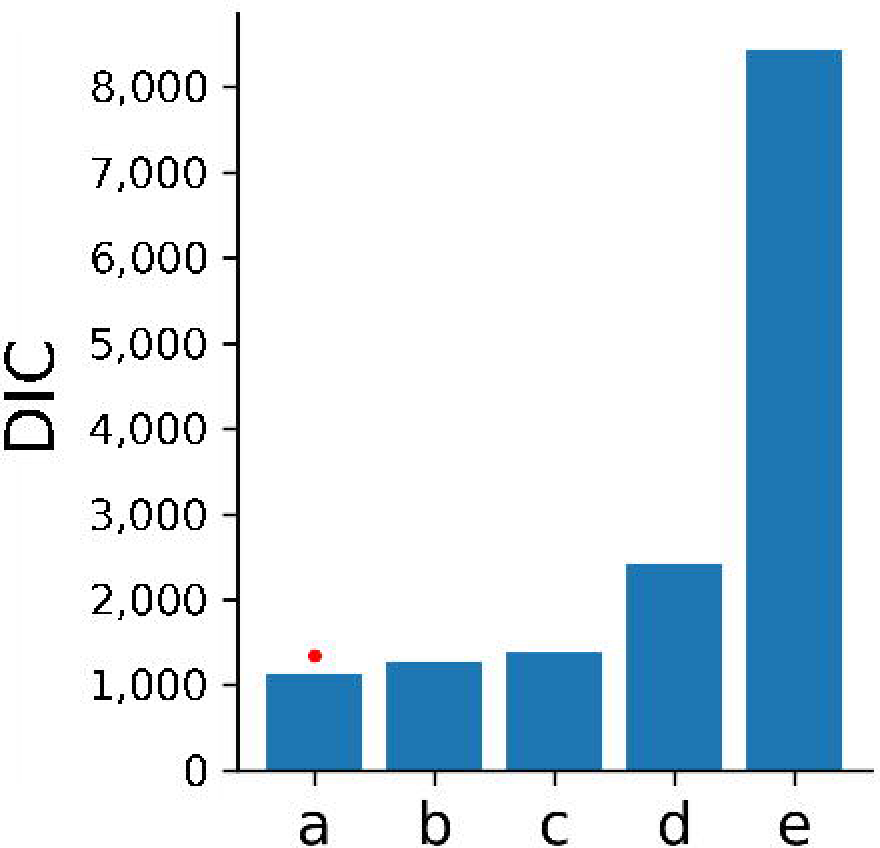
Model comparison. Description of the models (v = drift, a = decision boundary, t = non-decision time, z = bias; conditions: ses = session, diff = difficulty, const = constant across conditions):

a: v(ses*diff)_a(ses)_t(ses)_z(ses)

b: v(ses*diff)_a(ses)_t(ses)

c: v(ses*diff)_a(const)_t(ses)_z(ses)

d: v(ses)_a(ses)_t(ses)_z(ses)

e: v(ses*diff)_a(ses)_t(const)_z(ses)

The best performing model is marked with a red dot. Model a has the lowest DIC value and outperforms the closest model by ΔDIC > 100.

### Learning on and specificity of components of the decision-making process

To dissociate different aspects of the decision-making process and learning effects upon them, we use Hierarchical Drift Diffusion Models (HDDM) to relate accuracy to reaction times. To this end, we consider the data from Experiment 1 (joystick – saccade) in which the task was performed under time pressure (by means of a fixed response time window of 1 s) and without a delay phase; the DDM is most appropriate for such speeded reaction times [57]. We consider the parameters drift rate, decision boundary, bias, and non-decision time. To find the combination of model parameters which best explains our data, we fit several models and compare their DIC values (Fig. 6). The model which allows drift rate to vary across sessions and difficulty levels and decision boundary, non-decision time and bias to vary only across sessions provides the best fit to our data (model a). This model outperforms the closest model (b), which does not account for bias, by a difference in DIC >100, which is considered definitive evidence for the better model. According to Gelman-Rubin method, all parameters of model (a) converged with a value < 1.016 and there is no difference in convergence quality between sessions (Fig. S2). The Geweke method indicates that all group parameters (*n*=40) have an average *z*-score ∈ [-1.5, 1], and at worst, 70% of the values are within two standard deviations of zero (Fig. S3). All subject parameters (*n*=796) have an average *z*-score = ±2, which is within acceptable bounds. Only few parameters have 40-70% of the values within two standard deviations of zero. Also, the fit quality is not different across sessions in each of the four independent runs (Fig. S4). Together, this shows that our HDDMs converged well. Additionally, we verify the model through a posterior predictive comparison (Table S1), which demonstrates that the predicted accuracy and reaction time distributions match the empirical data.

#### Drift rate

Both when assessed on the group and on the subject level, we find that drift rate increases between the first and the last training session across difficulty levels (Fig. 7A, *p*_subj_(*v*_last_training_ > *v*_first_training_) = [0.95, 0.99, 0.99, 0.99, 0.99], see Table S2 for SD of subject values, *p*_group_(*v*_last_training_ > *v*_first_transfer_) = [1.0, 1.0, 1.0, 1.0, 1.0] for ±Δ_orientation_ = [0.31°, 0.56°, 1°, 1.77°, 3.16°], respectively). This is in keeping with previous reports that show that VPL increases the drift rate in various tasks [e.g., 33,34]. When we change the effector in the transfer session, drift rates drop, which is evident in the single subject and, especially, group fit of the HDDM (*p*_subj_(*v*_last_training_ > *v*_transfer_) = [0.79, 0.66, 0.85, 0.77, 0.86], see Table S2 for SD of subject values, *p*_group_(*v*_last_training_ > *v*_transfer_) = [0.98, 0.85, 0.99, 0.95, 0.99]). We do not find this difference in drift rate between effectors in the untrained control group (*p*_group_(*v*_hand_> *v*_eyes_) = [0.45, 0.23, 0.56, 0.26, 0.35], *p*_group_(Δ*v_Exp1_* > Δ*v_Control_*) = [0.85, 0.86, 0.81, 0.90, 0.92]). We note that statistical power of the between-groups comparison may be affected by higher variance in the drift estimation (see Fig. S5). In the control group, drift was conditioned on effectors and not sessions, which may lead to an increase in the variance of the MCMC samples. Nevertheless, the results overall suggest that learning effects on the rate of evidence accumulation depend on the effector.

**Figure 7:**
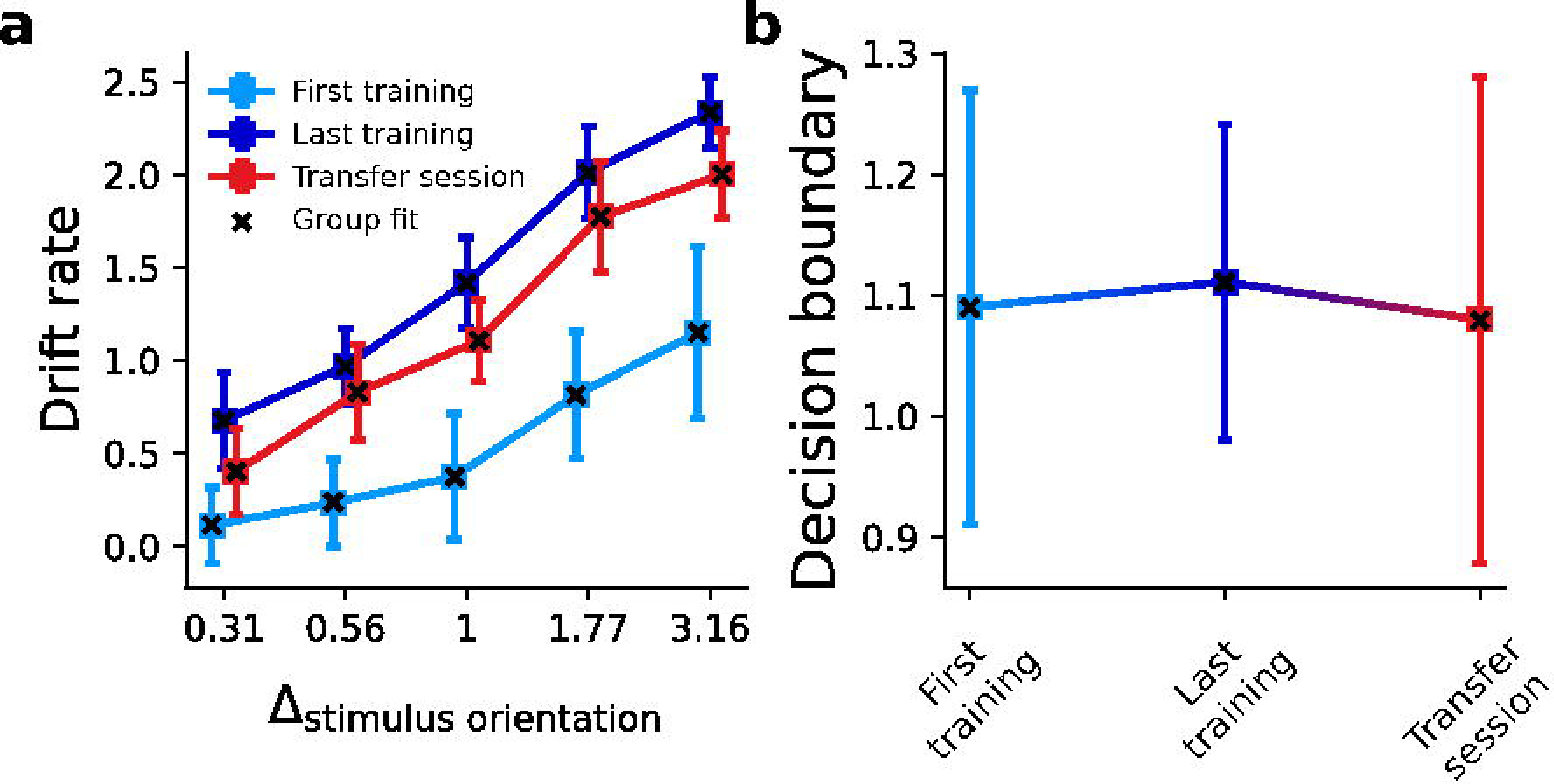
Hierarchical Drift Diffusion Model fit of drift rates and decision boundaries in Experiment 1. **A.** Drift rate (v) increases between the first and the last training session across orientations. When the effector changes in the transfer session, the learning-induced gains in drift rate drop. **B.** The separation between decision boundaries (a) does not change between the first, last training and transfer session. Squares signify the mean individual fits, errors bars ±2 standard deviations; crosses indicate the group fits. The discrepancy along the abscissa between sessions in A is intentional to increase visibility of error bars.

#### Decision boundaries

We find that the separation between decision boundaries corresponding to clockwise and counterclockwise choices does not change significantly between sessions (Fig. 7B, *p*_subj_(*a*_last_training_ > *a*_first_training_) = 0.58±0.35SD, *p*_group_(*a*_last_training_ > *a*_first_training_) = 0.73, *p*_subj_(*a*_last_training_ > *a*_transfer_) = 0.59±0.38SD, *p*_group_(*a*_last_training_ > *a*_transfer_) = 0.81). There is also no difference in decision boundaries between effectors in the untrained control group and the trained group (Fig. S6, *p*_group_(Δ*a_Exp1_* > Δ*a_Control_*) = 0.53).

#### Non-decision time

Non-decision time increases between the first training session and the last training session, as is clearly evident on the group level (Fig. 8A, *p*_group_(*t*_last_training_ > *t*_first_training_) = 0.99), but less consistently on the subject level (*p*_subj_(*t*_last_training_ > *t*_first_training_) = 0.78±0.39SD, see Table S2 for SD of subject values). Previous studies have also reported increases in non-decision time with learning [e.g., 35]. It is well established that non-decision time can be adjusted strategically, e.g., under different speed-accuracy trade-off instructions [58-60]. Hence, it is possible that subjects adjusted their motor execution time to stay within the response time window, i.e., to avoid too fast responses while their drift rate increases. Non-decision time decreases massively in the transfer session (*p*_subj_(*t*_last_training_ > *t*_transfer_) = 1.0±0.00SD, *p*_group_(*t*_last_training_ > *t*_transfer_) = 1.0, *p*_subj_(*t*_first_training_ > *t*_transfer_) = 0.87±0.30SD, *p*_group_(*t*_first_training_ > *t*_transfer_) = 1.0, without three extremely fast subjects *p*_subj_(*t*_first_training_ > *t*_transfer_)=1.0±0.00SD). The non-decision time in the untrained control group is also significantly smaller for eyes (*p*_group_(*t*_hand_ > *t*_eyes_) = 0.99) and the difference between the control and trained group is negligible (Fig. S7, *p*_group_(Δ*t_Exp1_* > Δ*t_Control_*) = 0.57). This effect can likely be attributed to generally shorter latencies of saccades as compared to hand movements [40,46].

#### Bias

We find a slight bias in clockwise orientation (Fig. 8B, *z*_subj_ = 0.471, *z*_group_ = 0.471) during the first training session, which disappears with the last training (*z*_subj_ = 0.505, *z*_group_ = 0.505; *p*_subj_(*z*_last_training_ > *z*_first_training_) = 0.77±0.29SD, *p*_group_(*z*_last_training_ > *z*_first_training_) = 0.99). Hence, subjects seem to optimally adjust their response bias in accordance with the task demands. During the transfer session, the bias is almost eliminated (*z*_subj_ = 0.493, *z*_group_ = 0.493; *p*_subj_(*z*_last_training_ > *z*_transfer_) = 0.59±0.26SD, *p*_group_(*z*_last_training_ > *z*_transfer_) = 0.79).

**Figure 8:**
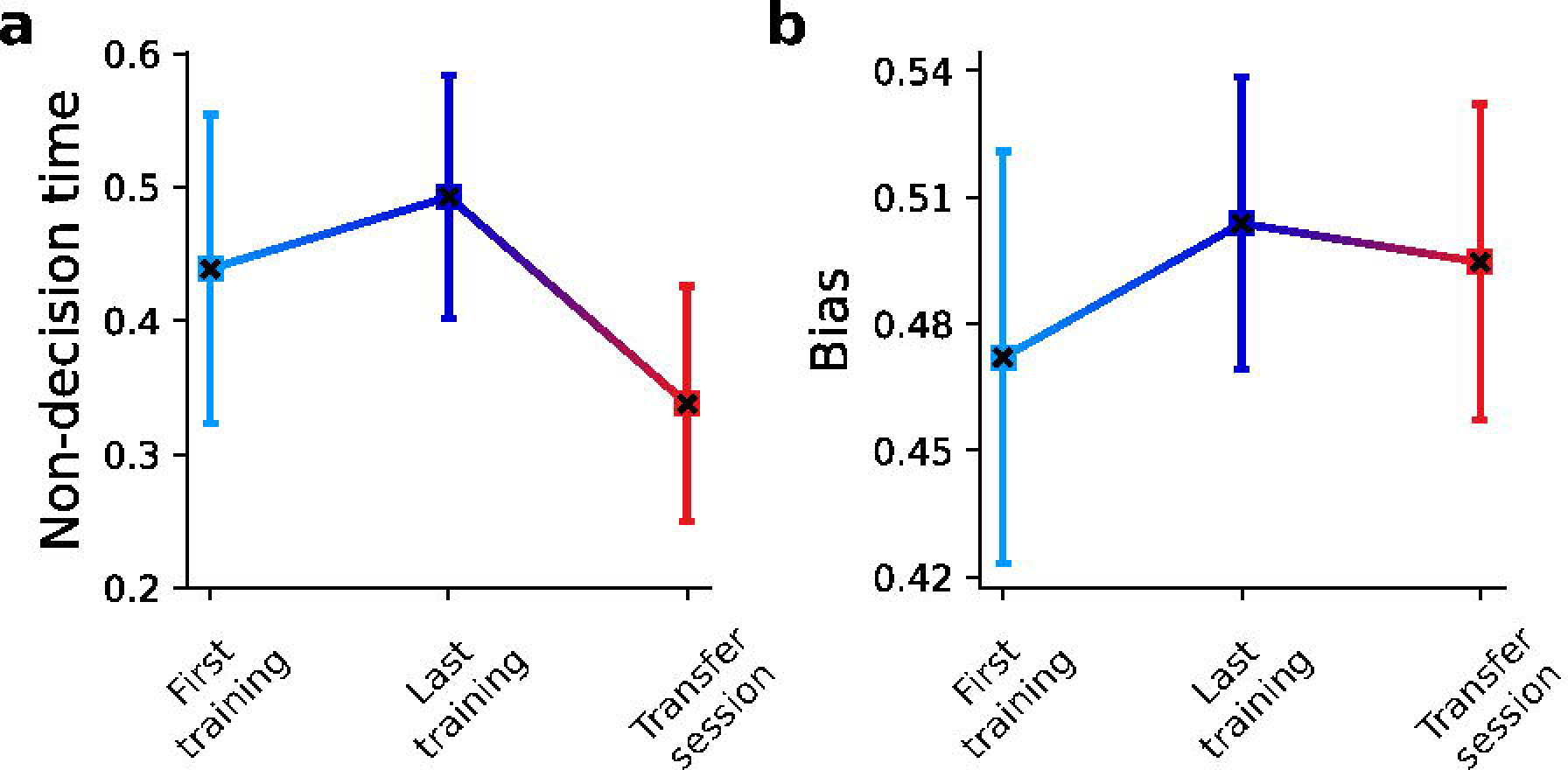
Hierarchical Drift Diffusion Model fit of non-decision times and biases in Experiment 1. **A.** Non-decision time (t) increases between the first and the last training session, and then decreases in the transfer session **B.** Subjects had a slight bias (z) in the clockwise direction during the first training session, which disappeared with the last training and remained low for the transfer session. Squares signify the mean individual fits, errors bars ±2 standard deviations; crosses indicate the group fits.

## Discussion

We find that VPL of orientation discrimination depends on the effector with which the task is trained. Analyses with the HDDM reveal that this dependency arises from effector specificity of a subcomponent of the decision-making process, namely the rate of evidence accumulation. Our results thus confirm that sensorimotor mapping is affected by VPL but suggests a new model on how and where this mapping may take place.

Previous models of VPL suggest either full or (little to) no transfer between effectors. Full transfer would arise if learning took place entirely in visual cortex [3-5], as originally conceived, or if sensorimotor mapping occurs between visual and decision making neurons, e.g., in parietal cortex, in the form of learning an abstract “rule” [25]. In the most drastic case, no transfer between effectors would be predicted if sensorimotor mapping would take place between visual and extremely effector-specific neurons in motor or parietal cortex [28] (but see below). In theory, learning abstract rules leads to excellent generalization capacity, which has for example been suggested to be the basis of visual training from action video games [61]. On the other hand, decomposing tasks into effector-specific components greatly simplifies reinforcement learning and effector-specific reward prediction errors supporting such learning have been found in humans using fMRI [62]. Effector specificity has been found in several other forms of learning involving sensorimotor mapping [e.g., 63], but generalization across effectors is also not uncommon [reviewed in 64]. Our results can be interpreted as an *effector-specific decision rule* (although neurophysiological data would of course be necessary to further support this interpretation). Here, sensorimotor mapping would occur between visual neurons and effector-specific integrators, e.g., in parietal cortex, similar to the *specific stimulus-response mapping model*.

However, the degree of specificity we find falls between the two previously reported extremes, as transfer is neither full nor none. Given the co-localization of saccade-specific, reach-specific, and effector-invariant motor neurons in parietal cortex [65,66], it is possible that there is overlap of the populations of neurons encoding motor plans for these two effectors. The degree of overlap would then determine the degree of transfer between the trained and untrained effectors in the transfer session. A possible reason for the colocalization of saccade- and reach-selective neurons in parietal cortex is the need for eye-hand coordination in many real-world tasks. Saccades may always be planned along limb movements for visual guidance, but not executed if fixation is required by the task [67]. Such consistent coactivation could also account for partial transfer results, as saccades would be trained along hand movements in an implicit fashion, perhaps through Hebbian plasticity.

In Experiment 2, we used a spatially-localized visually-guided pointing movement, as compared to the simple left/right joystick tilt response in Experiment 1, to match as well as possible the spatial contingencies of the saccade task. This was done to ensure that the finding of partial transfer in the first experiment was not due to very different spatial transformation between joystick movements and saccades but indeed dependent on the effector: there are considerable differences in parietal and occipital cortex activity associated with simple non-visual actions such as joystick tilt or a button press, compared to visually-guided actions where the effector (the hand, or a joystick cursor) is visible and needs to be directed to a spatial target [e.g., 68,69]. In line with this, lesions of parietal cortex influence spatially-contingent reach-and-point responses but do not affect arbitrary stimulus to simple non-spatial motor response transformations [70]. At the same, index finger pointing and saccades activate largely overlapping areas in parietal and prefrontal cortex [71], making the learning transfer question very relevant. Since we find significant SIs in both experiments as well as when we pool data across experiments, this indicates that VPL is indeed effector-specific to some degree; however, given that only Experiment 2 included a delay phase and that there was only a numerical but not a statistically significant drop in accuracy when the effector was changed suggests that further research is required to illuminate the role of spatial mapping in this form of specificity.

Classical VPL models in which learning takes place only in visual cortex predict full transfer and cannot easily explain partial specificity of learning effects to effectors. However, visual cortex activity has been shown to be differentially modulated by saccades versus reaches in humans using fMRI [67,72]. In rodents, there is evidence that motor effects in visual cortex increase information about the orientation of visual stimuli [reviewed in 73]. This suggests that interactions between visual and motor components of VPL could hypothetically already arise in early visual areas but still points to sensorimotor interactions as a relevant contributor to VPL.

Because neurons in parietal cortex have been shown to integrate visual information towards a decision bound, the parameters of the DDM are often interpreted to arise in parietal cortex as well [74]. This would fit with the known existence of effector-specific integrators and their colocalization in areas like LIP that form the basis of the newly suggested effector-specific but overlapping integrators model. An increased drift rate is usually interpreted as better sensory input to decision-making process [33,75]. However, this interpretation cannot fully account for the effector specificity of the drift rate parameter that we found, because the sensory input does not change when the effector changes. Rather, our results suggest that the drift rate (also) reflects the integration process itself, i.e., the action of the integrator. For example, it is possible that integrators in parietal cortex increase their sensitivity to sensory information, which could further contribute to a fast integration of information towards the decision bound.

Our results are in partial conflict with previous reports on full transfer [25] and no transfer between effectors [28,29]. As mentioned in the introduction, the latter results can conceivably be explained by the task-specificity of VPL (but we also note that they were obtained with a different experimental design than ours). Several possibilities could account for the discrepancy with the study by Awada and colleagues: the study by Awada and colleagues tested a much smaller sample of subjects (6 and 4 subjects in their Experiment 1 and 2, respectively), which may have been insufficient to reveal partial transfer effects. In fact, the results from Awada and colleagues’ Experiment 1 show a small trend towards partial transfer that did not reach statistical significance. However, it is also possible that differences in the stimuli and task account for the divergent results. In contrast to our study on orientation discrimination, Awada and colleagues investigated motion direction discrimination VPL. Motion direction is processed in dorsal stream areas, which are anatomically closer to parietal decision-making areas than neurons encoding orientation, which primarily reside in the ventral stream. Furthermore, parietal areas such as LIP contain intrinsically motion direction sensitive (i.e., sensory) neurons [76]. Both these factors could ease sensorimotor mapping to multiple colocalized effectors. Further studies with high statistical power explicitly comparing motion direction and orientation discrimination tasks will be able to resolve these alternative interpretations.

In addition to the effector-specific learning effects discussed above, there may have been concurrent, stimulus-unspecific motor learning. Yet, reaction times, which would be a straightforward marker of such effects, did not significantly decrease with training in Experiment 1, only Experiment 2 in our study. This difference between the two experiments is very likely due to the fact that the required movements in Experiment 1 were very simple and stereotyped, whereas the movements in Experiment 2, especially during the training phase with the touch screen, required more learning. General improvements in motor execution times would factor into the non-decision time parameter of the HDDM. To what extend general improvements in motor execution contribute to the effector-specific effects reported here is difficult to determine because the change in effector also entails a change in motor execution time. However, given that there was no difference between the experimental group undergoing training and the control group undergoing no training, it seems unlikely that there were such effects at play in Experiment 1, which may be due to the very simple and stereotypical movement that needed to be executed which is likely to be already highly overtrained. An additional form of learning that may factor into our results is general task learning irrespective of the stimulus and/or the effector. Task learning could be revealed by changing the task, not the effector, and previous studies indeed suggest that such learning takes place [e.g., 29]. It is possible that general task learning reduced the degree of effector specificity in our study. Dissociating the different components of plasticity involved in and/or concurrently occurring with visual perceptual learning remains an important target for future experiments. [e.g., 30]. It is possible that general task learning reduced the degree of effector specificity in our study. Dissociating the different components of plasticity involved in and/or concurrently occurring with visual perceptual learning remains an important target for future experiments. [e.g., 29]. It is possible that general task learning reduced the degree of effector specificity in our study. Dissociating the different components of plasticity involved in and/or concurrently occurring with visual perceptual learning remains an important target for future experiments.

Taken together, we find that VPL is not as specific, but also not as generalizable as previously thought. Taking into account different components of the sensorimotor arc revealed additional aspects of VPL beyond the traditional focus on (early) visual areas. Our results also suggest an expanded interpretation of the drift rate as reflecting not only the amount of accumulable information, but also the sensitivity of the integrator itself. Finally, in terms of application of VPL [77], where generalization is key, it may be beneficial to counteract overtraining by varying effectors. Given the new effector-specific learning rule model we propose, more effector variability during training may enable more generalization of training outcomes [also see 78,79].

## Acknowledgements

This project was supported by a seed fund from the Leibniz ScienceCampus ‘Primate Cognition’, Göttingen, Germany (to CMS and IK), and has received funding from the European Research Council (ERC) under the European Union’s Horizon 2020 research and innovation programme (Grant agreement No. 802482, to CMS). CMS is supported by the German Research Foundation Emmy Noether Program (SCHW1683/2-1). The funders had no role in study design, data collection and interpretation, decision to publish, or preparation of the manuscript. The authors declare no competing financial or non-financial interests.

## Author contributions

*Vladyslav Ivanov:* Methodology, Software, Formal analysis, Investigation, Writing - Original Draft, Visualization. *Giorgio L. Manenti:* Methodology, Formal analysis, Investigation, Data Curation, Writing - Original Draft, Visualization. *Sandrin S. Plewe:* Methodology, Investigation, Writing - Review & Editing. *Igor Kagan:* Conceptualization, Writing - Original Draft. *Caspar M. Schwiedrzik:* Conceptualization, Methodology, Writing - Original Draft, Supervision, Project administration, Funding acquisition.

## Data availability

The datasets generated and/or analyzed during the current study will be made publicly accessible upon acceptance of the manuscript in a peer-reviewed journal.

## Notes

### Competing Interest Statement

The authors have declared no competing interest.

### Summary of Updates

We have performed several new analyses of reaction times and accuracy data, especially concerning the factor task difficulty, that were requested during peer review. We have also added new figures and re-plotted existing figures in accordance with the peer review. The conclusions of the manuscript remain unchanged. We note that one of the co-authors has changed names.

