## Supplementary Material for "Decision-making processes in perceptual learning depend on effectors"

**Supplemental Material**

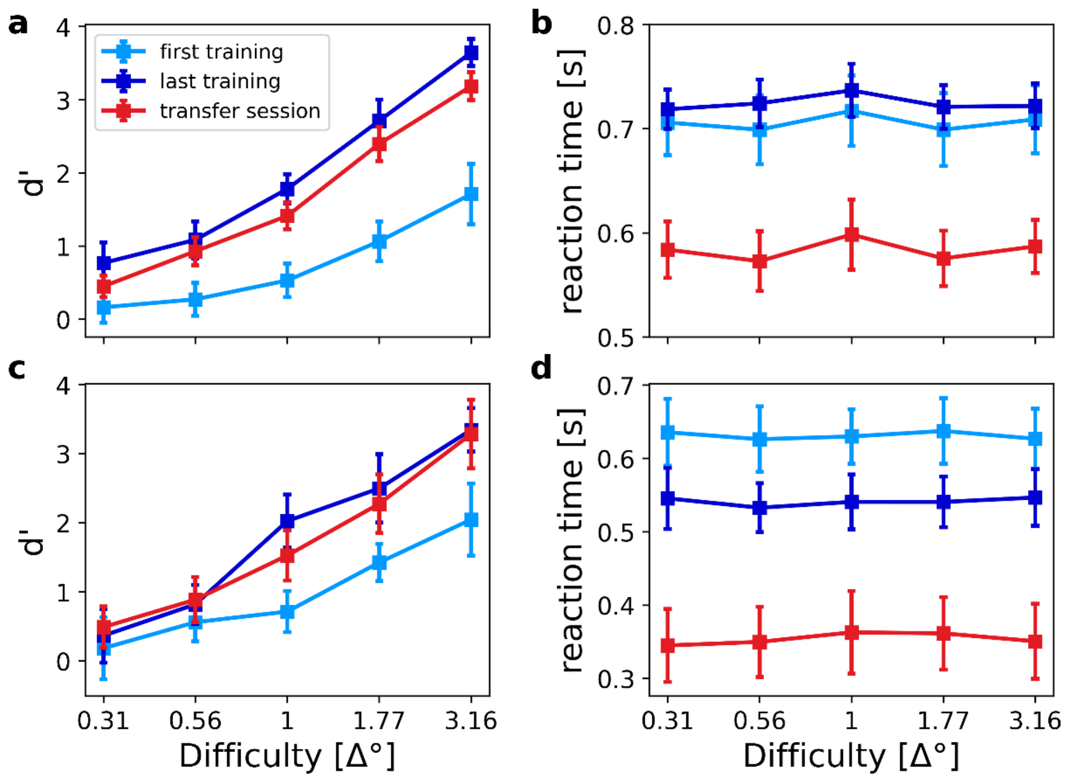


**Figure S1. D' and reaction time per difficulty level. A.** Mean d' and **B.** mean reaction time across subjects for Experiment 1. Here, we find no significant main effect of or interaction with difficulty in the reaction times when we compare the first to the last training session (rmANOVA, all $p>0.05$, all partial $\eta^2<0.11$). There is a small main effect of difficulty when we compare the last training session to the transfer session (rmANOVA, $F(4,72)=2.6468$, $p=0.0402$, partial $\eta^2=0.13$), which varies with a complex quartic trend over difficulty levels ($t(72)=2.899$, $p=0.005$). There is, however, no interaction with the effector change (rmANOVA, $F(4,72)=0.8918$, $p=0.4734$, partial $\eta^2=0.0472$), suggesting that the difficulty effect is not effector specific. **C.** Mean d' and **D.** mean reaction time across subjects for Experiment 2. We also find no significant main effect of or interaction with difficulty in the reaction times when we compare the first to the last training session (rmANOVA, all $p>0.4585$, all partial $\eta^2<0.0918$), replicating the results from Experiment 1. In this experiment, there is no main effect of difficulty or interaction with difficulty when we change the effector (rmANOVA, all $p>0.3631$, all partial $\eta^2<0.1105$). In all panels, difficulty is indicated as angular degrees of deviation from the reference stimulus (left side = hardest condition, right side = easiest condition). Error bars indicate the standard error of the mean, corrected for between-subject variability [1,2].

In the reaction time analyses reported here, we only considered correct responses. We additionally performed rmANOVAs of reactions times including the factor accuracy. A substantial number of subjects did not make errors in the easy conditions in some sessions. We replaced these missing values with the mean of the remaining subjects in the same condition.

Using this approach, in Experiment 1, comparing the first training session with the last training session, there no statistically significant main effect of difficulty ($F(4,72)=1.51$, $p=0.2082$, partial $\eta^2=0.07$), but a borderline significant interaction between difficulty and accuracy ($F(4,72)=1.91$, $p=0.1165$, partial $\eta^2=0.09$). Comparing the last training session with the transfer session, there is again no main effect of difficulty ($F(4,72)=1.79$, $p=0.1389$, partial $\eta^2=0.09$), and the interaction closest to statistical significance is the interaction between difficulty and accuracy ($F(4,72)=1.56$, $p=0.1934$, partial $\eta^2=0.07$). This suggests that there was possibly a weak modulation of reaction times by difficulty in this experiment, but that it was either too weak or too variable to reach customary significance thresholds.

In Experiment 2, comparing the first with the last training session, the main effect of difficulty approaches significance ($F(4,36)=2.14$, $p=0.0950$, partial $\eta^2=0.19$). Here, we do find a significant 2-way interaction between session and difficulty ($F(4,36)=4.48$, $p=0.0048$, partial $\eta^2=0.33$), and a significant interaction between accuracy and difficulty ($F(4,36)=2.91$, $p=0.0346$, partial $\eta^2=0.24$). This mainly reflects a trend for longer reaction times for easier orientation differences in the incorrect responses in the last training session. Comparing the last training session with the transfer session in Experiment 2, the main effect of difficulty is significant ($F(4,36)=3.74$, $p=0.0119$, partial $\eta^2=0.29$), as is the interaction between accuracy and difficulty ($F(4,36)=3.08$, $p=0.0278$, partial $\eta^2=0.25$). These effects reflect a similar trend across orientation difference as for the analysis of the first and last training session, especially when responses are incorrect.

**Table S1. Comparison of the observed and simulated data (parameter recovery).** We simulated data based 500 values taken from the posterior distribution of each parameter. We used the posterior predictive check function provided by the HDDM python library. Response = average boundary choice on (0 := clockwise, 1 := counter clockwise). RT ≔ reaction time, 10q ≔ $10^{th}$ quantile of the reaction time distribution, ub = upper decision boundary (counterclockwise), lb = lower boundary (clockwise). SEM = standard error from the mean, MSE = mean-squared error. The column “credible” indicates whether the simulated data based on the fitted model falls within the 95% credible interval.

| | Statistic | Observed | Simulated | | | | |
|---|---|---|---|---|---|---|---|
| | | mean | mean | std | SEM | MSE | credible |
| | **Response** | 0.467 | 0.490 | 0.288 | 0.000514 | 0.0838 | True |
| **RT** | **mean (ub)** | 0.690 | 0.712 | 0.125 | 0.000497 | 0.0163 | True |
| | **std (ub)** | 0.166 | 0.166 | 0.095 | 0.000000 | 0.0092 | True |
| | **10q (ub)** | 0.483 | 0.560 | 0.105 | 0.005892 | 0.0170 | True |
| | **30q (ub)** | 0.599 | 0.609 | 0.111 | 0.000093 | 0.0124 | True |
| | **50q (ub)** | 0.675 | 0.668 | 0.123 | 0.000038 | 0.0152 | True |
| | **70q (ub)** | 0.758 | 0.753 | 0.147 | 0.000022 | 0.0217 | True |
| | **90q (ub)** | 0.916 | 0.907 | 0.212 | 0.000092 | 0.0454 | True |
| | **mean (lb)** | -0.692 | -0.703 | 0.122 | 0.000131 | 0.0152 | True |
| | **std (lb)** | 0.168 | 0.166 | 0.094 | 0.000006 | 0.0089 | True |
| | **10q (lb)** | 0.483 | 0.552 | 0.103 | 0.004711 | 0.0153 | True |
| | **30q (lb)** | 0.591 | 0.600 | 0.108 | 0.000072 | 0.0118 | True |
| | **50q (lb)** | 0.675 | 0.658 | 0.120 | 0.000267 | 0.0148 | True |
| | **70q (lb)** | 0.766 | 0.743 | 0.144 | 0.000547 | 0.0213 | True |
| | **90q (lb)** | 0.916 | 0.898 | 0.209 | 0.000314 | 0.0441 | True |

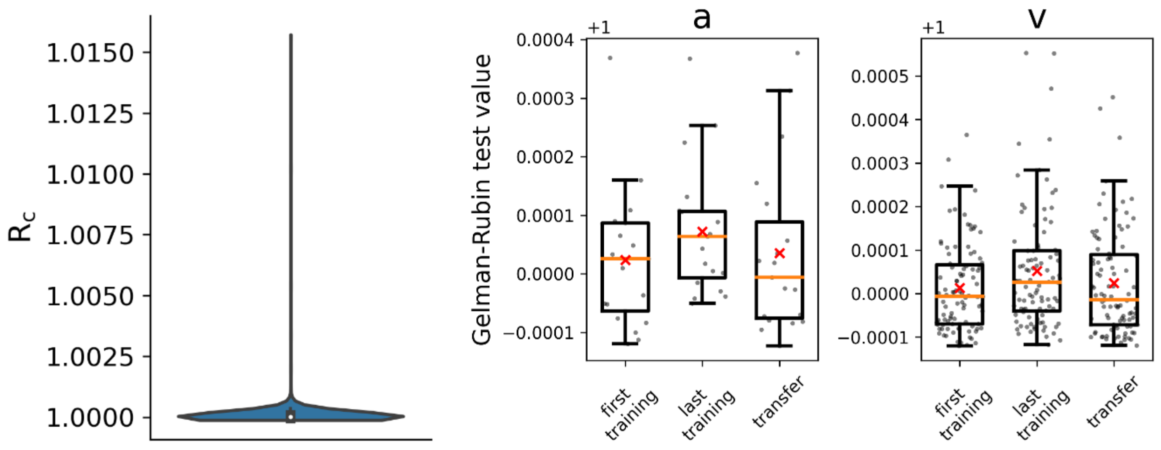


**Figure S2. Gelman-Rubin diagnostics. A.** Gelman-Rubin convergence test (number of runs = 4). **B.** Visualization of the convergence for the parameters per sessions. Orange line is median and the red cross is mean.

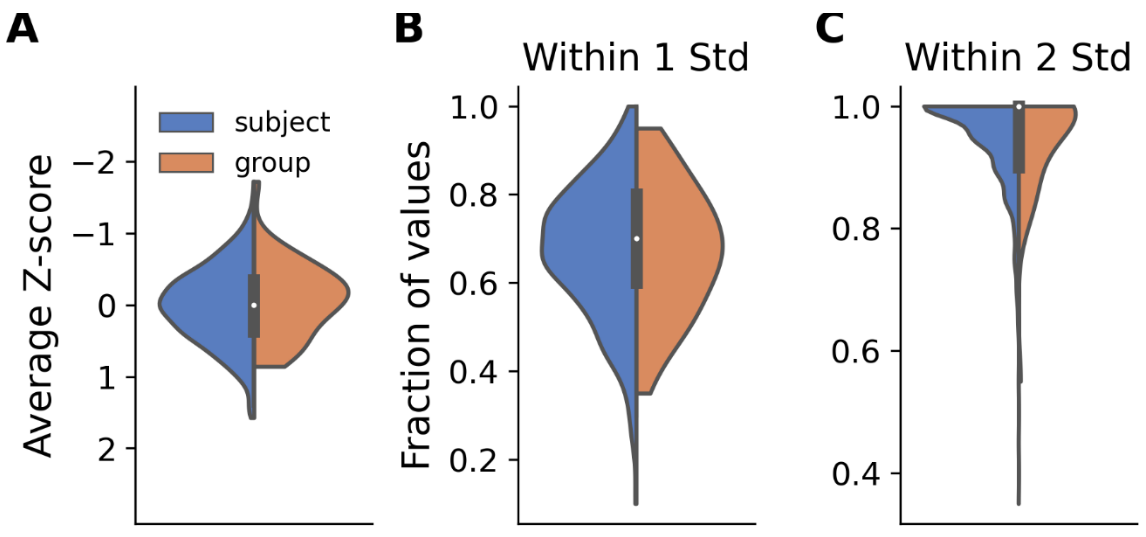


**Figure S3. Geweke convergence diagnostics.** ***A.*** Geweke convergence diagnostics for subject (n=798) and group (n=40) parameters. The distributions were calculated with a Gaussian kernel density estimator. **B-C.** The fraction of values within one or two standard deviations around zero across parameters.

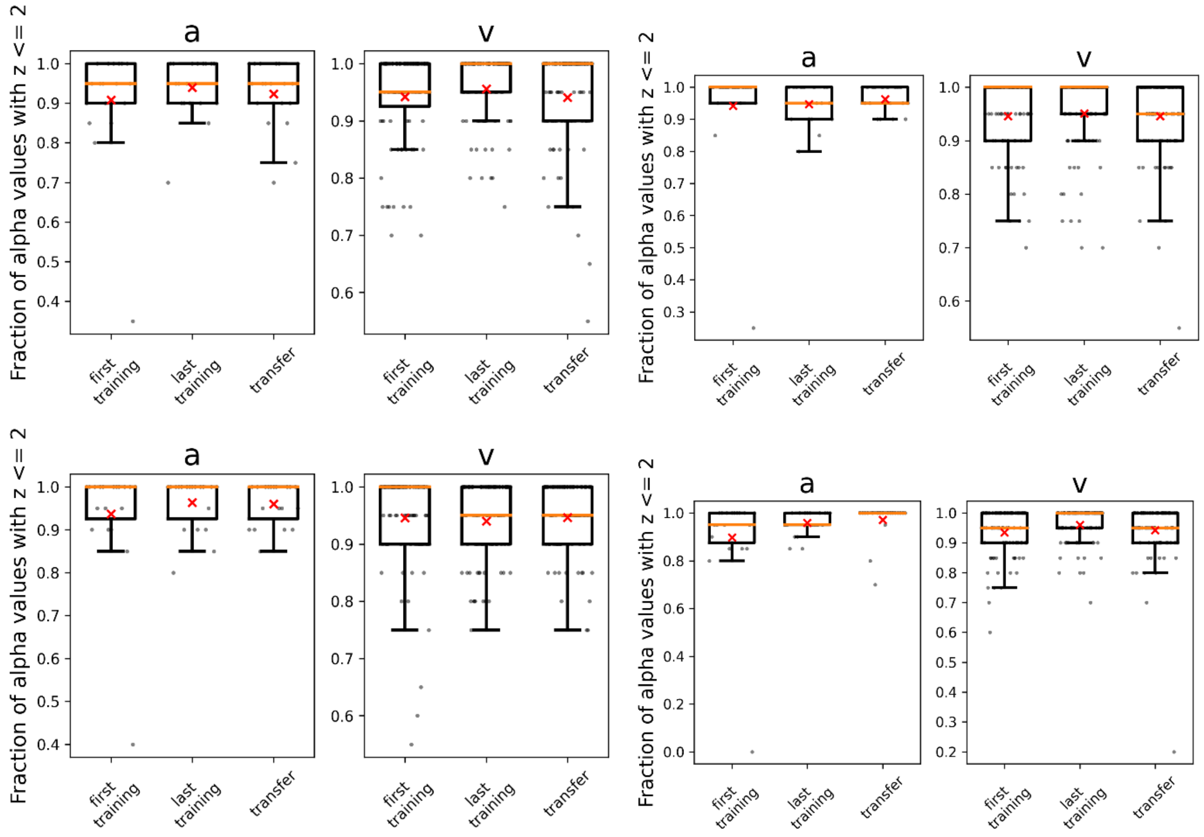


**Figure S4. Geweke convergence diagnostics across sessions and runs.** Each quadrant consisting of two plots with titles "a" and "v" represent independent runs. A - decision boundary, v – drift rate. Ordinate units are the same as in Fig. S3C.

**Table S2.** Comparison of HDDM conditions for the model with constant decision boundary across difficulties (Fig. 6, model *b*). $v_x$ = drift rate for *x* orientation, *a* = decision boundary, *t* = non-decision time, *z* = bias, $p_{4>1}$ = probability a random sample from session 4 is bigger than a random sample from session 1 (session 4 is the last training, etc.). The values in column "Subject" are the averages of the per subject comparison, SD = standard deviation.

| Parameter | $p_{\text{last training > first training}}$ | | $p_{\text{last training>transfer}}$ | |
|---|---|---|---|---|
| | **Subject** | **Group** | **Subject** | **Group** |
| $v_{0.31}$ | 0.95±0.10SD | 1.0 | 0.79±0.17SD | 0.98 |
| $v_{0.56}$ | 0.99±0.01SD | 1.0 | 0.66±0.18SD | 0.85 |
| $v_{1.0}$ | 0.99±0.00SD | 1.0 | 0.85±0.09SD | 0.99 |
| $v_{1.77}$ | 0.99±0.00SD | 1.0 | 0.77±0.16SD | 0.95 |
| $v_{3.16}$ | 0.99±0.00SD | 1.0 | 0.86±0.09SD | 0.99 |
| ***a*** | 0.59±0.35SD | 0.73 | 0.59±0.38SD | 0.82 |
| ***t*** | 0.78±0.39SD | 0.99 | 1.0±0.00SD | 1.0 |
| ***z*** | 0.77±0.29SD | 0.99 | 0.59±0.26SD | 0.79 |

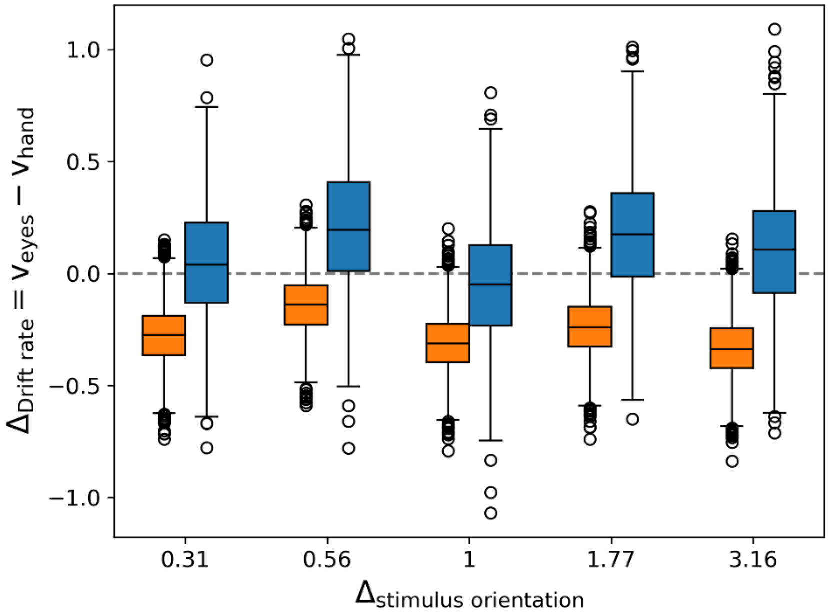


**Figure S5: Comparison of drift (group level parameters) difference between effectors in Experiment 1 and the control experiment.** The blue boxes represent the control experiment and the orange boxes Experiment 1. The x-axis indicates the difficulty levels as Δ° from the reference orientation (smallest Δ° = hardest condition) and y-axis the difference between the drift MCMC samples for both effectors ($\Delta v = v_{eyes} - v_{hand}$). A negative difference indicates that eyes performed worse than hand.

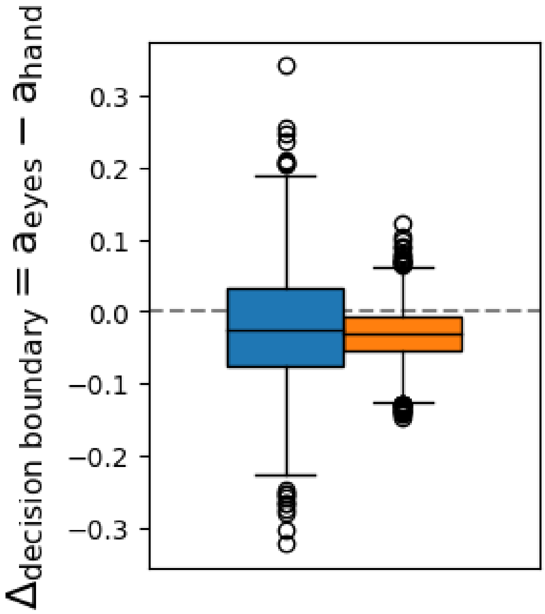


**Figure S6: Comparison of decision boundary (group level parameters) difference between effectors in Experiment 1 and the control experiment.** The blue boxes represent the control experiment and the orange boxes Experiment 1. The y-axis indicates the difference between the decision boundary MCMC samples for both effectors ($\Delta a = a_{eyes} - a_{hand}$). A negative difference indicates that eyes had a lower boundary.

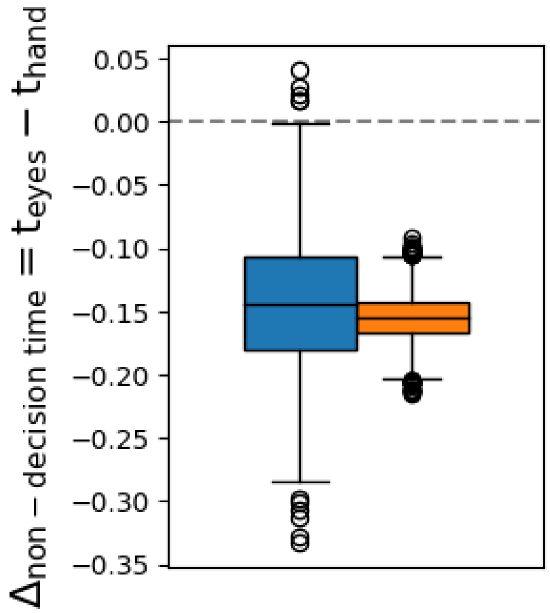


**Figure S7: Comparison of non-decision time (group level parameters) difference between effectors in Experiment 1 and the control experiment.** The blue boxes represent the control experiment and the orange boxes Experiment 1. The y-axis indicates the difference between the non-decision time MCMC samples for both effectors ($\Delta t = t_{eyes} - t_{hand}$). A negative difference indicates that eyes had lower reaction latency.

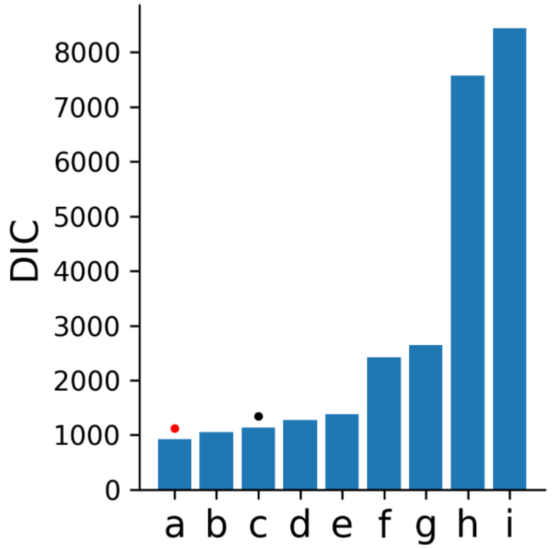


**Figure S8: Model comparison.** Description of the models (v = drift, a = decision boundary, t = non-decision time, z = bias; conditions: ses = session, diff = difficulty, const = constant across conditions):

**a:** v(ses*diff)_a(ses*diff)_t(ses)_z(ses)
**b:** v(ses*diff)_a(ses*diff)_t(ses)
**c:** v(ses*diff)_a(ses)_t(ses)_z(ses)
**d:** v(ses*diff)_a(ses)_t(ses)
**e:** v(ses*diff)_a(const)_t(ses)_z(ses)
**f:** v(ses)_a(ses)_t(ses)_z(ses)
**g:** v(ses)_a(ses*diff)_t(ses)_z(ses)
**h:** v(ses*diff)_a(ses*diff)_t(const)_z(ses)
**i:** v(ses*diff)_a(ses)_t(const)_z(ses).

The best performing model is marked with a red dot. Model **a** has the lowest DIC value and outperforms the closest model by ΔDIC > 100. The model allows decision boundary to vary also within sessions between difficulty levels. Model **c** is the model we used for the analysis in the main text (black dot). Its decision boundary varies only between sessions. Model **a** outperforms model **c** by by ΔDIC > 200.

**Table S3. Comparison of the observed and simulated data (parameter recovery) for the model with the variable decision boundary across sessions and difficulty levels.** We computed the same metric as in Table S1.

| | Statistic | Observed | Simulated | | | | |
|---|---|---|---|---|---|---|---|
| | | mean | mean | std | SEM | MSE | credible |
| | **Response** | 0.46771 | 0.49182 | 0.3042 | 0.00058 | 0.09312 | True |
| **RT** | **mean (ub)** | 0.69045 | 0.70813 | 0.12155 | 0.00031 | 0.01509 | True |
| | **std (ub)** | 0.16605 | 0.16253 | 0.09022 | 1E-05 | 0.00815 | True |
| | **10q (ub)** | 0.48336 | 0.55884 | 0.10809 | 0.0057 | 0.01738 | True |
| | **30q (ub)** | 0.59992 | 0.60734 | 0.11154 | 6E-05 | 0.0125 | True |
| | **50q (ub)** | 0.67501 | 0.66519 | 0.12094 | 0.0001 | 0.01472 | True |
| | **70q (ub)** | 0.75839 | 0.7477 | 0.14074 | 0.00011 | 0.01992 | True |
| | **90q (ub)** | 0.91668 | 0.89856 | 0.19846 | 0.00033 | 0.03972 | True |
| | **mean (lb)** | -0.6923 | -0.70084 | 0.11966 | 7E-05 | 0.01439 | True |
| | **std (lb)** | 0.16899 | 0.1628 | 0.08918 | 4E-05 | 0.00799 | True |
| | **10q (lb)** | 0.48337 | 0.5522 | 0.10706 | 0.00474 | 0.0162 | True |
| | **30q (lb)** | 0.59171 | 0.59943 | 0.11023 | 6E-05 | 0.01221 | True |
| | **50q (lb)** | 0.67501 | 0.65663 | 0.11919 | 0.00034 | 0.01454 | True |
| | **70q (lb)** | 0.76667 | 0.73935 | 0.13849 | 0.00075 | 0.01993 | True |
| | **90q (lb)** | 0.91668 | 0.89211 | 0.19579 | 0.0006 | 0.03894 | True |

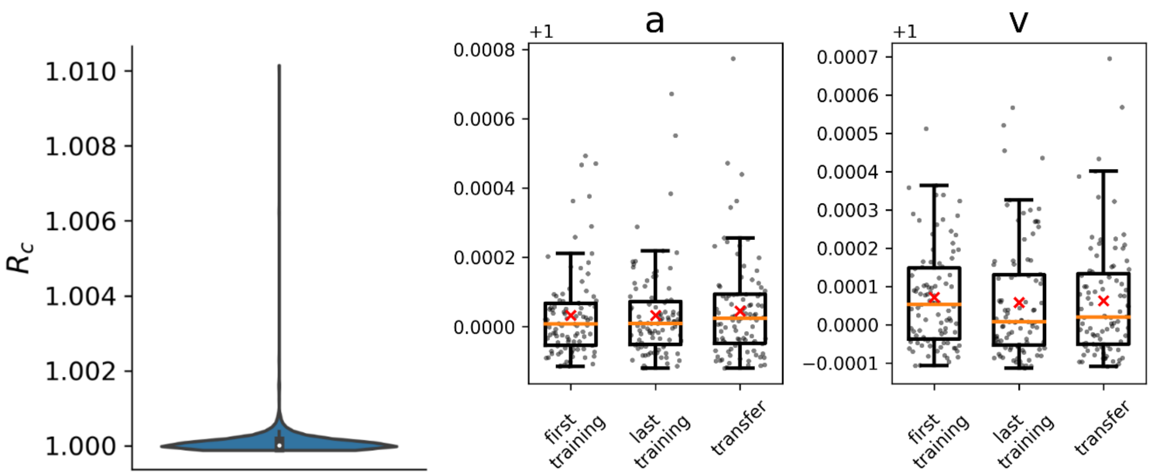


**Figure S9. Gelman-Rubin diagnostics. A.** Gelman-Rubin convergence test (number of runs = 4). **B.** Visualization of the convergence for the parameters per sessions. Orange line is median and the red cross is mean.

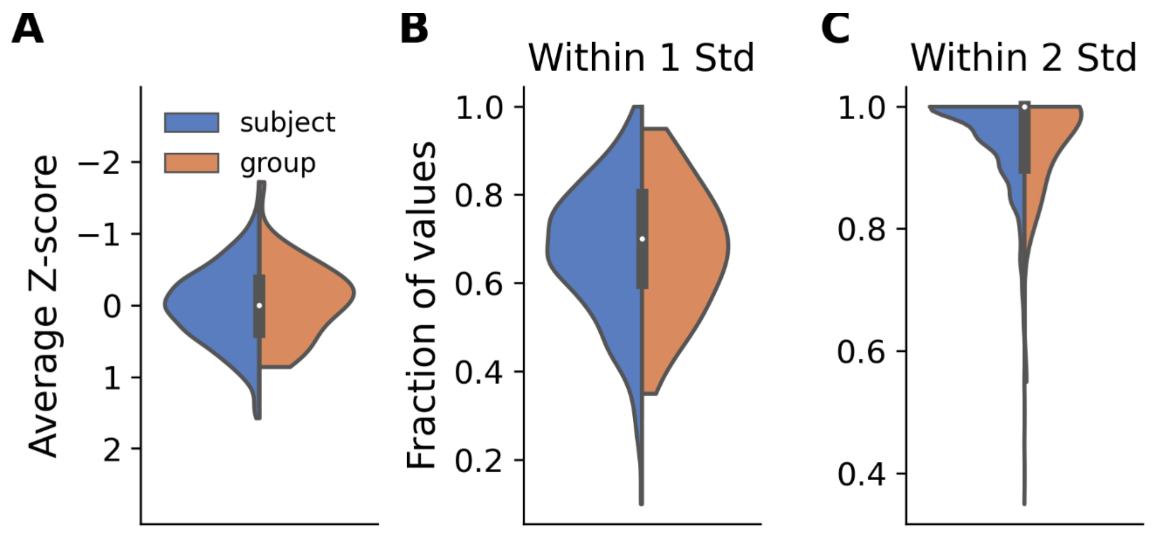


**Figure S10. Geweke convergence diagnostics.** ***A.*** Geweke convergence diagnostics for subject (n=1138) and group (n=64) parameters. The distributions were calculated with a Gaussian kernel density estimator. **B-C.** The fraction of values within one or two standard deviations around zero across parameters.

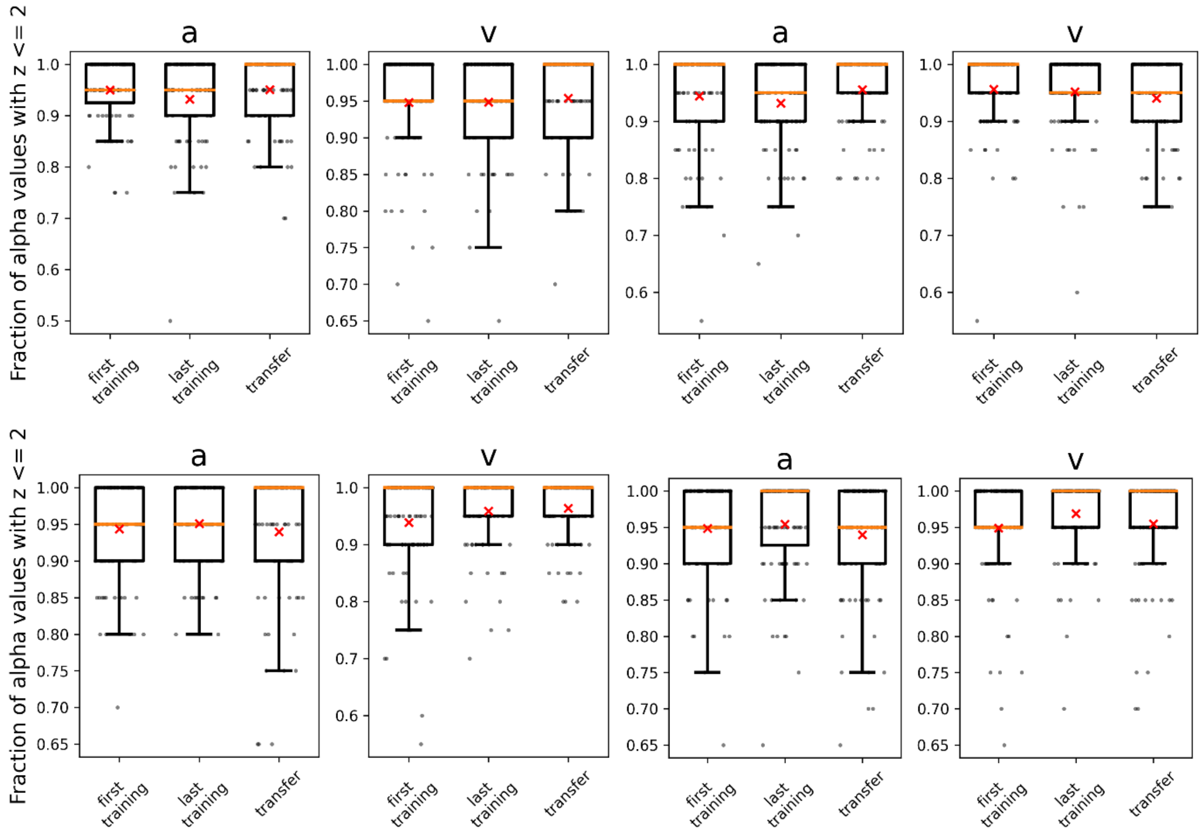


**Figure S11. Geweke convergence diagnostics across sessions and runs.** Each quadrant consisting of two plots with titles "a" and "v" represent independent runs. A - decision boundary, v – drift rate. Ordinate units are the same as in Fig. S3C.

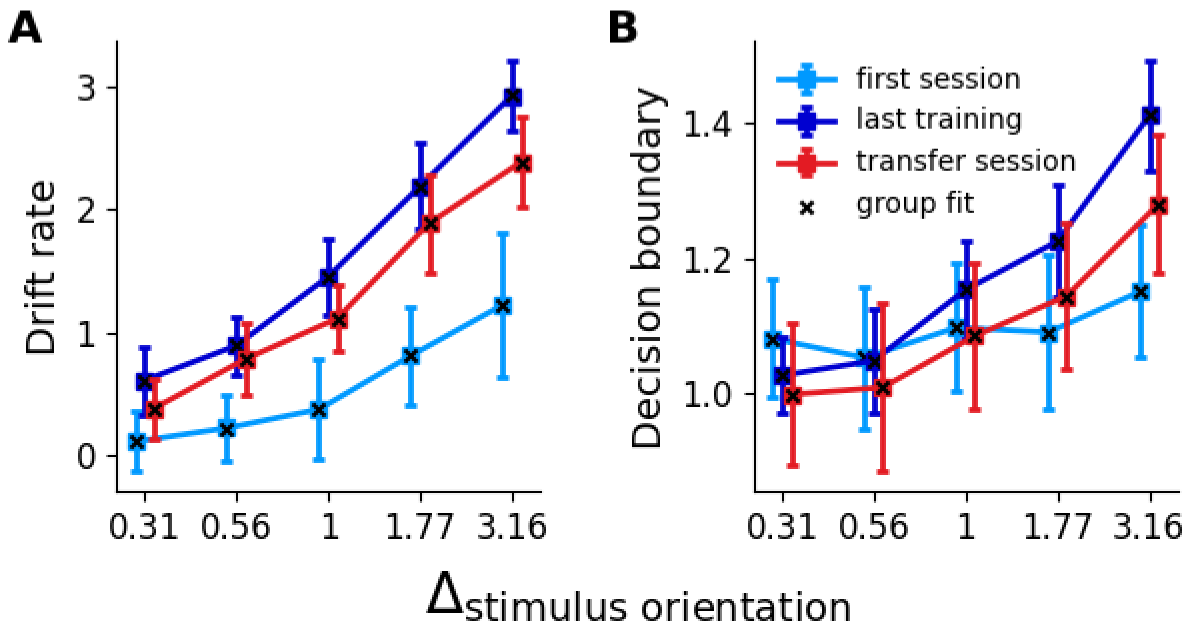


**Figure S12: Hierarchical Drift Diffusion Model fit of drift rates and decision boundaries. A.** Drift rate (v) increases between the first and the last training session across orientations. When the effector changes in the transfer session, the learning-induced gains in drift rate drop. **B.** The separation between decision boundaries (a) increases between the first and last training session for the easier orientations and decreases or stays the same for the most challenging conditions. Learning effects drop when the effector changes in the transfer session. Squares signify the mean individual fits, errors bars ±2 standard deviations; crosses indicate the group fits. The discrepancy along the abscissa between sessions is intentional to increase visibility of data points.

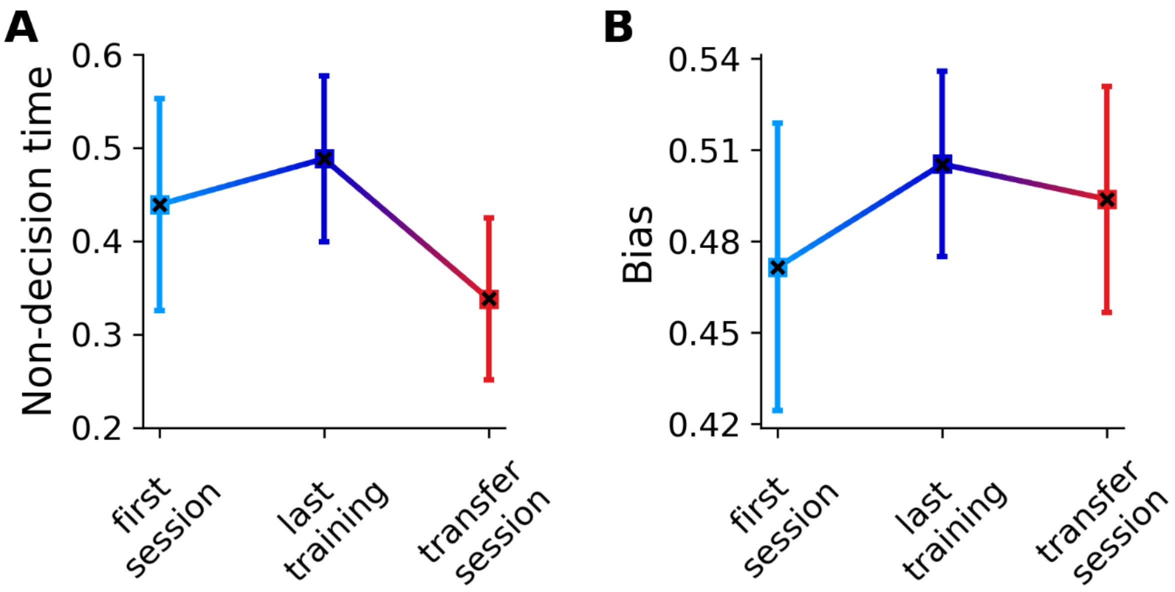


**Figure S13: Hierarchical Drift Diffusion Model fit of non-decision times and biases. A.** Non-decision time (t) increases between the first and the last training session, and then decreases in the transfer session. **B.** Subjects had a slight bias (z) in the clockwise direction during the first training session, which disappeared with the last training and remained low for the transfer session. Squares signify the mean individual fits, errors bars ±2 standard deviations; crosses indicate the group fits.

**Table S4.** Comparison of HDDM conditions. $v_x$ = drift rate for $x$ orientation, $a$ = decision boundary for $x$ orientation, $t$ = non-decision time, $z$ = bias, $p_{4>1}$ = probability a random sample from session 4 is bigger than a random sample from session 1 (session 4 is the last training, etc.). The values in column "Subject" are the averages of the per subject comparison, SD = standard deviation.

| Para-meter | $p_{last\ training > first\ training}$ | | $p_{last\ training>transfer}$ | |
|---|---|---|---|---|
| | **Subject** | **Group** | **Subject** | **Group** |
| $v_{0.31}$ | 0.92±0.13SD | 0.99 | 0.73±0.18SD | 0.95 |
| $v_{0.56}$ | 0.97±0.02SD | 1.0 | 0.61±0.19SD | 0.78 |
| $v_{1.0}$ | 0.99±0.00SD | 1.0 | 0.84±0.10SD | 0.99 |
| $v_{1.77}$ | 0.99±0.00SD | 1.0 | 0.78±0.19SD | 0.97 |
| $v_{3.16}$ | 1.0±0.00SD | 1.0 | 0.91±0.09SD | 0.99 |
| $a_{0.31}$ | 0.25±0.16SD | 0.05 | 0.60±0.27SD | 0.80 |
| $a_{0.56}$ | 0.45±0.26SD | 0.44 | 0.63±0.28SD | 0.87 |
| $a_{1.0}$ | 0.70±0.20SD | 0.93 | 0.74±0.17SD | 0.96 |
| $a_{1.77}$ | 0.86±0.16SD | 0.99 | 0.76±0.17SD | 0.97 |
| $a_{3.16}$ | 0.94±0.02SD | 1.0 | 0.83±0.14SD | 0.99 |
| $t$ | 0.77±0.38SD | 0.99 | 1.0±0.00SD | 1.0 |
| $z$ | 0.79±0.27SD | 0.99 | 0.63±0.24SD | 0.85 |

**Model with variable decision boundaries across session and difficulty levels**

In the model with variable decision bounds per difficulty level, results in terms of drift rate closely correspond to the model with a single difficulty level for all difficulty levels. However, we also find that the separation between decision boundaries corresponding to clockwise and counterclockwise choices increases between the first and the last training session, most clearly and consistently in the easier conditions, but decreases or stays the same for the two most challenging conditions (Fig. S12B, $p_{subj}(a_{last\_training} > a_{first\_training})$ = [0.25, 0.46, 0.72, 0.89, 0.99], see Table S4 for subject SD values, $p_{group}(a_{last\_training} > a_{first\_training})$ = [0.05, 0.44, 0.93, 0.99, 1.0] ] for $\pm\Delta_{orientation}$ = [0.31°, 0.56°, 1°, 1.77°, 3.16°], respectively). The separation is reduced when we change the effector in the transfer session ($p_{subj}(a_{last\_training} > a_{transfer})$ = [0.60, 0.63, 0.74, 0.76, 0.83], see Table S4 for subject SD values, $p_{group}(a_{last\_training} > a_{transfer})$ = [0.8, 0.87, 0.96, 0.97, 0.99]). The difference in decision boundary between effectors in the untrained control group and the trained group is weak ($p_{group}(\Delta a_{Exp1} > \Delta a_{Control})$ = [0.41, 0.55, 0.65, 0.76, 0.79]). However, within the control group, the difference between effectors is insignificant ($p_{group}(a_{hand} > a_{eyes})$ = [0.74, 0.64, 0.65, 0.56, 0.67]) and the difference within effectors between groups indicates the presence of learning for the easier conditions ($p(a_{eyes\_exp1} - a_{eyes\_control})$ = [0.44, 0.41, 0.85, 0.94, 0.94]). Hence, like the drift rate, learning effects on the decision boundary depend on the effector.

**Strategic adjustment of decision boundary to drift**

We observe the emergence of a correlation between drift rate (*v*) and decision boundary (*a*) throughout training (Spearman correlation $\bar{\rho}_{first\_training}$=0.511, $p$=0.380; $\bar{\rho}_{last\_training}$=0.947, $p$=0.021; $t(18)$=5.88, $p$<0.001, $g$=1.70, see Table S5). This correlation remains significant during the transfer session (Spearman correlation $\bar{\rho}_{transfer}$=0.947, $p$=0.021; $t(18)$<0.001, $p$>0.999, $g\approx0$). We can relate this finding to the well-known speed-accuracy trade-off. The subjects, apparently, preferred speed over accuracy for more challenging conditions, and accuracy over speed for easier conditions. They lowered or stabilized *a* even after the improvement of *v* for the two most difficult orientations (see Fig. S12A-B). Given the low drift rates, the emphasis on speed reduced the probability of exceeding the response time limit (1.25s after the stimulus onset) and, hence, the overall number of trial attempts. For the three least demanding conditions, subjects strategically used the additional time afforded by an increase in the drift rate to sample additional information towards higher accuracy instead of saving time on task by decreasing their reaction times through tighter decision bounds. To quantify the interaction between *v* and *a*, we fit the first-degree polynomial to the data per subject and session, where *a* depends on *v* over difficulty (orientation) levels. Thus, the slope of the linear fit reflects how *a* changes with growing *v*, and the offset is *a* when *v* equals zero. Throughout the training, the slope and offset converge on higher and lower values, respectively (Fig. S14). This is evident from a significant difference in the *v*-*a* relationship between the first and last training session (2D minimum energy test, see below, $e = 2.391$, $p = 0.001$). During the transfer session, only the slope changes ($t(189) = 4.28$, $p = 0.0004$), while the offset is not different from the last training session ($t(18) = -0.77$, $p = 0.44$). The slight change in the slope can be explained by a non-linear interaction between *v* and *a*. Accordingly, the difference between the final training session and the transfer session is statistically significant in 2D ($e = 0.165$, $p = 0.04$); however, not in comparison with the penultimate training session ($e = 0.040$, $p = 0.65$). Considering the overall faster reaction times (by ca. 150 ms) during the transfer session, subjects could apparently compensate for the reduced *v* by raising *a*, which would be reflected in higher offset values.

To investigate the relationship between drift rate and boundary separation, we use a 2D minimum energy test [3,4], as implemented by B. Lau (https://github.com/brian-lau/multdist). This test assesses via permutations whether two multidimensional samples are drawn from the same distribution.

**Table S5.** Pearson (blue cells) and Spearman (orange cells) correlation between drift rate and decision boundary across orientation (difficulty) levels per session. The values in column "Subject" are the averages of per subject. SD = standard deviation.

| Session | Subject | | Group | |
|---|---|---|---|---|
| | coefficient | p value | coefficient | p value |
| **1** | 0.622±0.338SD | 0.250±0.220SD | 0.847 | 0.069 |
| **2** | 0.916±0.055SD | 0.032±0.031SD | 0.961 | 0.009 |
| **3** | 0.919±0.047SD | 0.030±0.025SD | 0.937 | 0.018 |
| **4** | 0.967±0.028SD | 0.008±0.012SD | 0.985 | 0.002 |
| **Transfer** | 0.929±0.054SD | 0.026±0.028SD | 0.967 | 0.007 |
| **1** | 0.510±0.349SD | 0.381±0.306SD | 0.8 | 0.1 |
| **2** | 0.811±0.159SD | 0.119±0.115SD | 0.9 | 0.037 |
| **3** | 0.761±0.146SD | 0.151±0.115SD | 0.6 | 0.28 |
| **4** | 0.947±0.061SD | 0.021±0.027SD | 1 | < 0.001 |
| **Transfer** | 0.947±0.061SD | 0.021±0.027SD | 1 | < 0.001 |

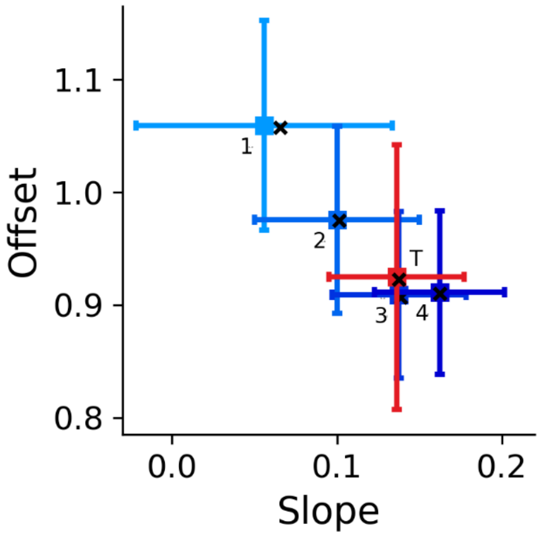


**Figure S14: Relationship between drift rate and decision boundary.** Slope and offset of a linear fit of drift rates against decision boundaries across orientations. Throughout the training sessions, the slope and the offset stabilize at higher and lower values, respectively. This is evidenced by a significant difference in the *a*-*v* relationship between the first and last training session (2D minimum energy test, $e$=2.391, $p$=0.001). During the transfer session, only the slope changes ($t(18)$=4.28, $p$=0.0004), while the offset remains the same as in the last training session ($t(18)$=-0.77, $p$=0.44). Colored error bars display the estimates based on the per-subject parameters (mean ±2 SDs), the black crosses show the estimates based on group parameters. Numbers indicate training sessions (1-4), transfer session is red and marked with T.

**Supplemental discussion**

When considering a model with variable decision bounds per difficulty level, we find that apart from the drift rate, VPL also has an effect on the decision bound, and that this effect is also (partially) effector-specific. Trainees seem to adjust their decision bound such that faster integration is in a way counteracted by a more conservative decision bound. This speed-accuracy policy results in higher accuracy because more information can be sampled but forgoes the opportunity to save time on task by reaching a decision earlier on the basis of a higher drift rate alone. Previous studies have reported a decreased decision bound [5] and less variability in the decision bound parameter [6] with VPL. However, differences in task design and instructions, e.g., an emphasis of speed over accuracy, may explain this discrepancy.

Our model with decision bounds that varied with the difficulty level of the task outperformed all other models. Usually, however, DDMs are instead fitted with a single decision bound that spans all difficulty levels and is thought to reflect a strategic criterion to optimize speed versus accuracy. Balci et al. [7] and Evans & Brown [8] suggest that with sufficient practice, subjects can learn to adjust their decision bound to difficulty levels. This is indeed in line with our results, as decision bounds are initially, i.e., in the first session, almost identical across difficulty levels, but differentiate only in later sessions (Fig. S14), speaking to a learning effect. Given this and that the model with decision bounds per difficulty level substantially outperformed the model with a single, fixed decision bound per session (Fig. S8), subject may have developed a sense of time to exploit the available response time window at maximum. Interestingly, the adjustment of $a$ to $v$ could be transferred to the new effector, suggesting that task strategies are not effector specific. However, we also note that the effector specificity of the drift rate parameter fully holds in the model with a single decision bound across difficulty levels that only varies between training sessions (Fig. 7 in main text).

Neurally, narrower decision bounds correlate with an increase in activity at the outset of the integration process in area LIP [9]. Since many neurons in LIP are effector specific, it is possible that the same neurons that may account for the learning effects and the specificity of the drift rate also account for the effects on the decision bound. Our model would thus predict a reduction in spontaneous choice-predictive activity in areas like LIP [10] with VPL. Alternatively, a central decision bound circuit, e.g., in the basal ganglia [11], where correlates of VPL have also been found [12], could account for our results.
